# Lifespan-extending downregulation of insulin signalling reduces germline mutation load

**DOI:** 10.64898/2025.12.05.692572

**Authors:** Elizabeth M. L. Duxbury, Alice M. Godden, Jean-Charles de Coriolis, Hanne Carlsson, Simone Immler, Alexei A. Maklakov

## Abstract

Reduced insulin/IGF-1 signalling (IIS) robustly extends lifespan and enhances somatic stress resistance across taxa, yet its consequences for germline genome integrity remain unclear. Here we combine multigenerational mutation accumulation with whole-genome sequencing in *C. elegans* to test whether adulthood-only IIS downregulation can simultaneously promote somatic maintenance and limit germline mutational burden. We reduced IIS by adult-onset *daf-2* RNAi in wild-type and heritable RNAi-deficient (*hrde-1*) backgrounds, allowing either spontaneous or UV-induced germline mutations to accumulate over multiple generations. In wild-type animals, reduced IIS lowered germline single-nucleotide mutation rates by up to ∼50% and prevented the UV-induced elevation in mutation rate, without detectable costs to fecundity or lineage persistence. By contrast, in *hrde-1* mutants the same intervention increased both point mutations and transposable-element–driven insertions under UV exposure, accelerating lineage extinction. Thus, the genome-protective effect of reduced IIS critically requires the germline nuclear Argonaute HRDE-1, which mediates small-RNA–guided epigenetic silencing. Functional annotation of germline variants revealed enrichment in pathways linked to development, cellular maintenance and conserved longevity regulators, including IIS and mTOR, and identified high-impact mutations in genes with human orthologs implicated in neurodegeneration and cancer. Our findings show that IIS can coordinate somatic and germline maintenance in concert, rather than in competition, through an HRDE-1–dependent epigenetic pathway. This work positions nutrient-sensing IIS as a central regulator of germline genome stability and suggests that IIS downregulation can reduce germline mutation load while extending lifespan, with broad implications for biogerontology and evolutionary biology.

## 1 Introduction

Ageing is profoundly influenced by evolutionarily conserved nutrient-sensing pathways, among which insulin/insulin-like growth factor-1 signalling (IIS) has emerged as a central regulator of lifespan and late-life health. Genetic or environmental reductions in IIS extend lifespan and enhance somatic stress resistance in *C. elegans* and across diverse taxa, including flies and mice, while modulation of IGF-1 signalling is associated with variation in human longevity (Kenyon, 2010; Murphy & Hu, 2013; Regan et al., 2020; Biglou et al., 2021). IIS integrates information about nutritional status to coordinate growth, metabolism, and stress responses, in part via the FOXO transcription factor DAF-16 and other downstream effectors (Murphy & Hu, 2013; Biglou et al., 2021). Adult-onset reductions in IIS in *C. elegans* can extend lifespan without detectable costs to reproduction and can even improve offspring fitness, highlighting the importance of age-specific IIS regulation for organismal ageing (Dillin et al., 2002; Lind et al., 2019; 2021; Sultanova et al. 2025). However, despite the central role of IIS in somatic ageing, it remains unclear whether and how IIS influences germline genome integrity and mutation load across generations. Addressing this question is essential for understanding how systemic ageing pathways shape both somatic healthspan and intergenerational genomic stability.

High-fidelity repair of the germline genome and proteome is costly, and the lower limit of mutation rate is sometimes suggested to result from resource allocation trade-offs (Kimura, 1967; Sniegowski et al., 2000; Furió et al., 2005; Agarwal & Wang, 2008). For example, individual-level trade-offs between germline and somatic maintenance, or germline maintenance and reproduction can shape the evolution of mutation rates (Maklakov & Immler, 2016; Berger et al., 2017; Chen et al., 2020; Lind et al., 2024). In line with this “expensive germline” hypothesis, germline removal in *Danio rerio* zebrafish led to faster somatic repair and reduced apoptosis following ionising irradiation (Chen et al., 2020). In *C. elegans*, germline ablation results in increased somatic maintenance and lifespan (Hsin and Kenyon 1999; Arantes-Oliveira et al. 2002; Antebi 2013; Ermolaeva et al. 2013; Shemesh et al. 2013) and germline reduction following nutritional stress is also associated with increased lifespan (Thondamal et al. 2014). However, germline removal in killifish (Moses et al., 2024) and *Caenorhabditis remanei* nematodes (Lind et al., 2024) improved only male, but not female, lifespan. Together, these results do not exclude the possibility that germline removal is perceived by the organism as germline damage leading to upregulation of somatic maintenance to ensure long-term survival and reproduction in a stressful environment without direct reallocation of resources. Indeed, DNA damage to germ cells from exogeneous and endogenous sources improves somatic resilience to stress via innate immune peptide signalling in *C. elegans* (Ermolaeva et al., 2013). Overall, while theoretically the costs of high-fidelity germline maintenance may constrain the evolution of germline mutation rates (GMR), direct evidence for such trade-offs is limited.

Genetic interventions in *C. elegans* and across several other species show that reducing insulin/IGF-1 signalling (IIS) has widespread somatic benefits-extending lifespan, improving somatic maintenance and stress resistance (Kenyon, 2010; Murphy & Hu, 2013; Biglou et al., 2021), such as to prolonged high temperature, oxidative stress (Lithgow et al., 1995; Honda & Honda, 1999; Holzenberger et al., 2003; Kenyon, 2010) and pathogen infection (Garsin et al., 2003; Evans et al., 2008; Duxbury et al., 2024). Confining the reduction in IIS to adulthood, by feeding RNA interference (RNAi) to downregulate the expression of the *daf-2* insulin receptor homolog in *C. elegans* from the onset adulthood, did not lead to costs for development, or offspring number (Dillin et al., 2002; Lind et al., 2019; 2021) and instead improved offspring fitness (Lind et al., 2019). Recent work showed that adulthood *daf-2* RNAi in *C. elegans* also improved lineage survival under multigenerational spontaneous or UV-induced mutation accumulation (MA) and even improved the fitness of offspring assayed after 20 generations of UV-induced MA (Duxbury et al., 2022). Therefore, we hypothesized that reduced adulthood IIS may also lower GMR.

Here, we tested the potential role of IIS in regulating germline single-nucleotide mutations and transposable-element (TE)-driven insertions, under both spontaneous and UV-induced multigenerational germline mutation accumulation, using whole genome sequencing of *C. elegans* MA lines. We find that adulthood-only downregulation of IIS (rIIS) reduces GMRs and that germline Argonaute, *hrde-1* was critical to mediate the effects. We show that rIIS in *hrde-1* mutant MA lines led to increased GMRs and TE-driven germline insertions under UV-induced mutagenesis. Together, our results show that insulin signalling plays an important role in regulating GMR and TE activity and that multigenerational protection under rIIS requires functional inheritance of RNAi.

## 2 Results and Discussion

### 2.1 Germline mutation rates are modulated by insulin signalling

To determine the effects of reduced insulin signalling (rIIS) during adulthood on germline mutation rates (GMRs) in *C. elegans*, we used our established replicated mutation accumulation (MA) lines in which mutations were allowed to accumulate for 40 generations, either spontaneously (n=400 MA lines) or were induced with UV irradiation (n=400) (Duxbury et al., 2022). We used feeding RNAi downregulation of the insulin receptor, *daf-2*, to reduce IIS during adulthood, in half of the spontaneous and UV-irradiated MA lines and ran all four experimental treatments across wild-type N2 and *hrde-1* mutant backgrounds (n=100 MA lines per treatment, per genetic background, across the eight MA treatment combinations described; see Duxbury et al., 2022). Intergenerational effects of *daf-2* RNAi treatment are discussed in Supplementary Material. The *hrde-1* mutant background was used to determine the potential role of heritable RNAi-mediated silencing in maintaining germline genome integrity under MA (Buckley et al., 2012; Spracklin et al., 2017). GMRs were quantified via whole genome sequencing in a subset of MA lines per treatment (Table 1) and calculated as the number of line-specific, heterozygous, single nucleotide polymorphisms (SNPs) per MA line, per generation, scaled by the number of callable genomic sites (see Methods for full details).

**Table 1.** Summary of mean and standard error for transition/transversion (Ts/Tv) ratios, sample sizes and number of MA generations at which germline mutation rates were quantified, across MA lines, per treatment.

| **Treatment** | **Mean**<br>**(standard error)**<br>**Ts/Tv** | **Number of**<br>**MA lines** | **Number of**<br>**MA generations** |
| --- | --- | --- | --- |
| **N2 + ev** | 0.607<br>(0.054) | 14 | 35 |
| **N2 + *daf-2* RNAi** | 0.618<br>(0.042) | 15 | 35 |
| **N2 + ev + UV** | 0.666<br>(0.033) | 11 | 25 |
| **N2 + *daf-2* RNAi + UV** | 0.624<br>(0.056) | 18 | 25 |
| ***hrde-1* mutant + ev** | 0.601<br>(0.029) | 25 | 20 |
| ***hrde-1* mutant + *daf-2* RNAi** | 0.547<br>(0.029) | 24 | 20 |
| ***hrde-1* mutant + ev + UV** | 0.586<br>(0.048) | 22 | 10 |
| ***hrde-1* mutant + *daf-2* RNAi + UV** | 0.603<br>(0.044) | 23 | 6 |

GMRs ranged over three-fold across the eight MA treatments, from 3.1 to 11.9 x 10^-8^ (Figure 1; Table 1). UV irradiation significantly increased GMRs across both wild-type N2 and *hrde-1* mutant genetic backgrounds (Gaussian GLM, overall UV effect: *t*=10.775, *p* < 0.001). The extent of the effect of UV irradiation on mutation rates depended on both the genetic background and the RNAi treatment (Gaussian GLM, UV x genetic background: *t*=-6.595, *p* <0.001; UV x RNAi: *t*=-3.915, *p*<0.001; UV x RNAi x genetic background: *t*=3.514, *p*<0.001; Figure 1; Table 1; Supplementary Tables 1 and 2). Interestingly, for N2 MA lines on *daf-2* RNAi, there was no increase in GMRs under UV irradiation after 25 generations of MA, compared with spontaneous GMRs after 35 generations of MA (Gaussian GLM, UV: *t*=0.948, *p*=0.351), unlike the effect of UV irradiation for N2 MA lines on the empty vector (ev) control, strongly suggesting that rIIS protected against elevated UV irradiation-induced mutagenesis. While absolute germline mutation rates vary across *C. elegans* MA studies (Meier et al. 2014; 2018; Konrad et al. 2019; Rajaei et al., 2021), perhaps in part due to between-study variation in the number of MA generations, the filtering of either heterozygous or homozygous mutations, or the *E. coli* strain used as food; our main focus was to determine the relative difference in germline mutation rates between rIIS and control treatments.

**Figure 1.**
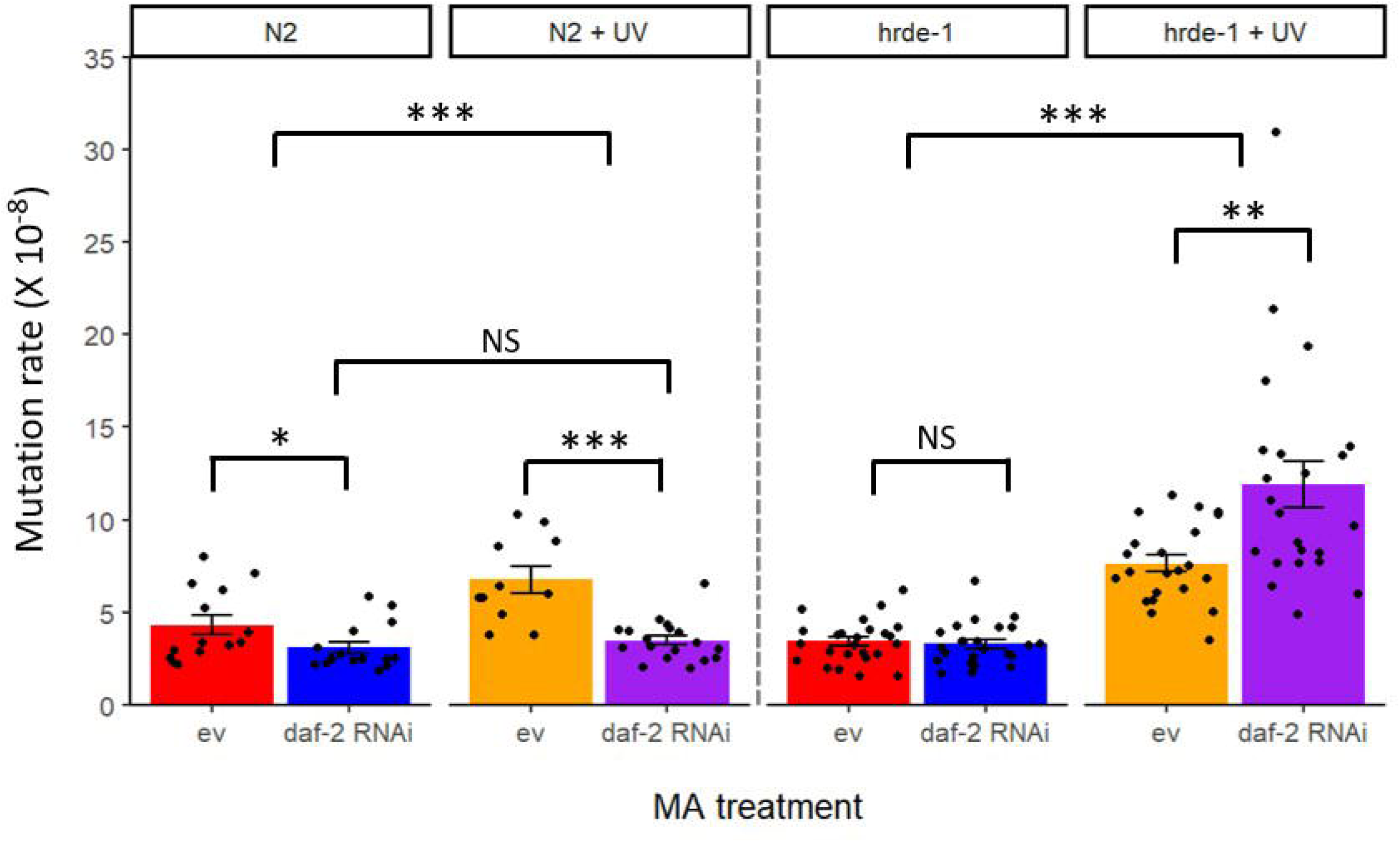
Heterozygous point mutation rates of *C. elegans* mutation accumulation (MA) lines across two genetic backgrounds: N2 wild type and *hrde-1* mutant and four MA treatments: empty vector (ev) control (red), reduced insulin/IGF-1 signalling via adulthood *daf-2* RNAi (blue), ev + UV irradiation (orange) or *daf-2* RNAi + UV (purple). Bars represent mean ± standard error and individual points represent the line-specific mutation rates, per generation, per callable genomic site for individual MA lines. Statistical significance: NS, non-significant; * p<0.05; ** p<0.01; *** p<0.001.

Reducing IIS lowered GMR and the number of mutations in N2 wildtype animals - most dramatically under UV-induced MA (Gaussian GLM for mutation rate: RNAi, t=5.153, p<0.001; Negative Binomial GLM for number of mutations: RNAi, *z*=5.665, *p*<0.001), but also when mutations accumulated spontaneously across generations (Negative Binomial GLM for number of mutations: RNAi, *z*=2.288, *p*=0.0221; although non-significant for mutation rate: RNAi, *t*=1.987, *p*=0.0571; RNAi x UV, *t*=2.362, *p*=0.0218; Figure 1; Table 1; Supplementary Tables 1-3). Mutation rates were positively associated with the extinction trajectories of the MA lines, such that MA lines from treatments with higher mutation rates also went extinct faster (Duxbury et al., 2022).

In contrast to N2 wildtype, *hrde-1* mutants under UV-induced MA, had a considerably increased mutation rate on *daf-2* RNAi, relative to *hrde-1* mutants under UV-induced MA on the ev control (Gaussian GLM for mutation rate: *t*=-3.189, *p*=0.00266; Figure 1; Table 1; Supplementary Table 1), which likely contributed to their rapid extinction (Duxbury et al., 2022). Under spontaneous MA, *daf-2* RNAi did not affect the mutation rate nor number of mutations for *hrde-1* mutants, relative to *hrde-1* mutant ev controls at generation 20 (Gaussian GLM: RNAi, *t*=0.435, *p*=0.665; RNAi x UV, *t*=-3.326, *p*=0.001; Negative Binomial GLM for number of mutations, RNAi, z=0.472, p=0.637; Figure 1; Supplementary Tables 2 & 3) and neither did their extinction trajectories differ (Duxbury et al., 2022). Mutation rates and extinction trajectories under spontaneous MA were similar between *hrde-1* mutants at generation 20 and N2 spontaneous MA lines on *daf-2* RNAi at generation 35 (Figure 1; Duxbury et al., 2022).

Overall, functional *hrde-1* was required for *daf-2* RNAi-mediated germline protection only when SNP mutation rates were elevated under UV-irradiation but had no apparent effect on spontaneous SNP mutation rates after 20 generations of MA (Figure 1). Spontaneous mutations may have accumulated non-linearly across MA generations in *hrde-1* mutants (as discussed in previous *C. elegans* MA work: Rajaei et al., 2021), such that mutation rate differences only became apparent in later generations beyond generation 20. Indeed, GMRs are supported by the minimal extinction seen in the first 20 generations of spontaneous MA for *hrde-1* mutants for both rIIS and control, compared to the rapid extinction of *hrde-1* mutants under UV-induced MA, accelerated further when combined with *daf-2* RNAi (Duxbury et al., 2022).

### 2.2 Mutational spectra were largely unaffected by insulin signalling or UV irradiation

To determine the nature of the germline mutational load between MA treatments and gain insight into possible mechanisms underlying the genomic responses to rIIS, we examined the broad- and fine-scale mutational spectra of SNPs. Transition (Ts; A:T->G:C, G:C->A:T) to transversion (Tv; A:T->C:G, G:C->T:A, A:T->T:A, G:C->C:G) ratios are expected to be 1:2 (Ts/Tv =0.5) if substitution rates are equal and occur randomly between nucleotides. Ts/Tv ratio did not significantly differ between MA treatments (Supplementary Table 4) but were all significantly greater than the random expectation of 0.5, except for non-irradiated *hrde-1* mutants on *daf-2* RNAi (range: 0.55-0.67; Table 1; Supplementary Table 5). Ts/Tv ratios were in the range of previous laboratory MA studies (Rajaei et al., 2021). These results indicate no significant effect of either rIIS or UV irradiation on the broad scale mutational spectrum of transitions and transversions (Supplementary Figure 1), but potentially enhanced purifying selection against transversions than transitions across all of the MA lines, as is seen in some wild isolates (Babbitt and Cotter 2011; Guo et al. 2017), although the exact mechanisms are unclear.

Under rIIS, N2 UV-irradiated MA lines, had a significantly greater proportion of A:T->C:G transversions (Supplementary Figure 2, Supplementary Table 6). MA treatments also significantly differed in the proportion of SNPs that were G:C->A:T transitions, but with no significant effect of rIIS specifically (Supplementary Table 6). Transversions were most commonly T to G or A to C across the MA lines (Supplementary Figure 2). While previous work found G:C->A:T transitions to be over-represented in MA lines relative to standing nucleotide diversity (Denver et al. 2009,2012; Konrad et al., 2019), perhaps indicative of elevated oxidative stress (Krašovec et al. 2017; Suzuki and Kamiya 2017; Poetsch et al. 2018; but see Rajaei et al., 2021), further work is required to determine why A:T->C:G transversions would be overrepresented under rIIS in N2 UV-irradiated MA lines, even though their overall germline mutational burden was lower than respective controls (Figure 1).

### 2.3 Elevated transposable element insertion rates in *hrde-1* mutants were mediated by IIS and UV irradiation

Since we found that rIIS, UV irradiation and genetic background (N2 versus *hrde-1* mutant) individually and jointly modulated germline single nucleotide point mutation rates in the MA lines, we next investigated MA treatment effects on transposable-element (TE)-induced insertion mutations. The aim was to test whether germline TE insertion rates, like germline SNP mutation rates were also elevated under UV-induced MA but lowered under rIIS and dependent on functional *hrde-1*.

TEs are estimated to comprise approximately 12% of the *C. elegans* genome and while several empirical studies suggest that many remain inactive, DNA transposons appear to be most actively transcribed (Bessereau et al., 2016; Sturm et al., 2023). Early work reported DNA transposons as the sole TE-driven contributor to spontaneous germline insertion mutations in *C. elegans* (Moerman & Waterson, 1984; Eide & Anderson, 1985), although *C. elegans* retrotransposon expression may only be detectable at certain temperatures and ages (Dennis et al., 2012). Furthermore, previous work has shown TEs to be activated under UV irradiation in *Drosophila simulans* fruit flies (Jardim et al., 2015) and at high temperatures in *C. elegans* (Sturm et al., 2023), *D. simulans* (Jardim et al., 2015) and *Arabidopsis* (Sun et al., 2020).

For this study we only considered TE insertions that were novel to the MA lines and neither found in the founder population from which the MA lines were initiated (for the respective N2 or *hrde-1* mutant genetic background) nor in the *C. elegans* reference genome. As such, filtered TE insertions were expected to have been induced by the MA treatments directly and presumed active, as opposed to the latent TEs already present in the genome prior to MA.

We found that genetic background had the strongest effect on MA line-specific TE insertion rates (Gaussian GLM, overall genetic background effect: *t*=-5.773, *p*<0.001). TE insertions rates were significantly higher in the *hrde-1* mutant background than in N2 wild types, for all MA treatments and especially for *hrde-1* mutants under UV-induced MA (Gaussian GLM, RNAi x UV x genetic background: *t*=2.899, *p*=0.004; UV x genetic background: *t*=-5.853, *p*<0.01; Figure 2; Supplementary Table 7). Strikingly, rIIS led to further increased TE insertion rates in UV-irradiated *hrde-1* mutant MA lines, relative to respective ev controls (Gaussian GLM, RNAi: *t*=-3.454, *p*=0.001; Figure 2). Under spontaneous MA, rIIS did not significantly reduce overall TE insertion rates for *hrde-1* mutants, relative to ev controls (Gaussian GLM, RNAi: *t*=0.981, *p*=0.331), as had been seen for the respective germline SNP mutation rates (Figure 1). Curiously, in N2 wildtype MA lines, neither rIIS nor UV irradiation had a significant effect on overall TE insertion rates (Gaussian GLM, RNAi: *t*=0.254, *p*=0.801, UV: *t*=0.838, *p*=0.405, RNAi x UV: *t*=0.651, *p*=0.518; Supplementary Table 7), which were all considerably lower than for the *hrde-1* mutant background (Figure 2).

**Figure 2.**
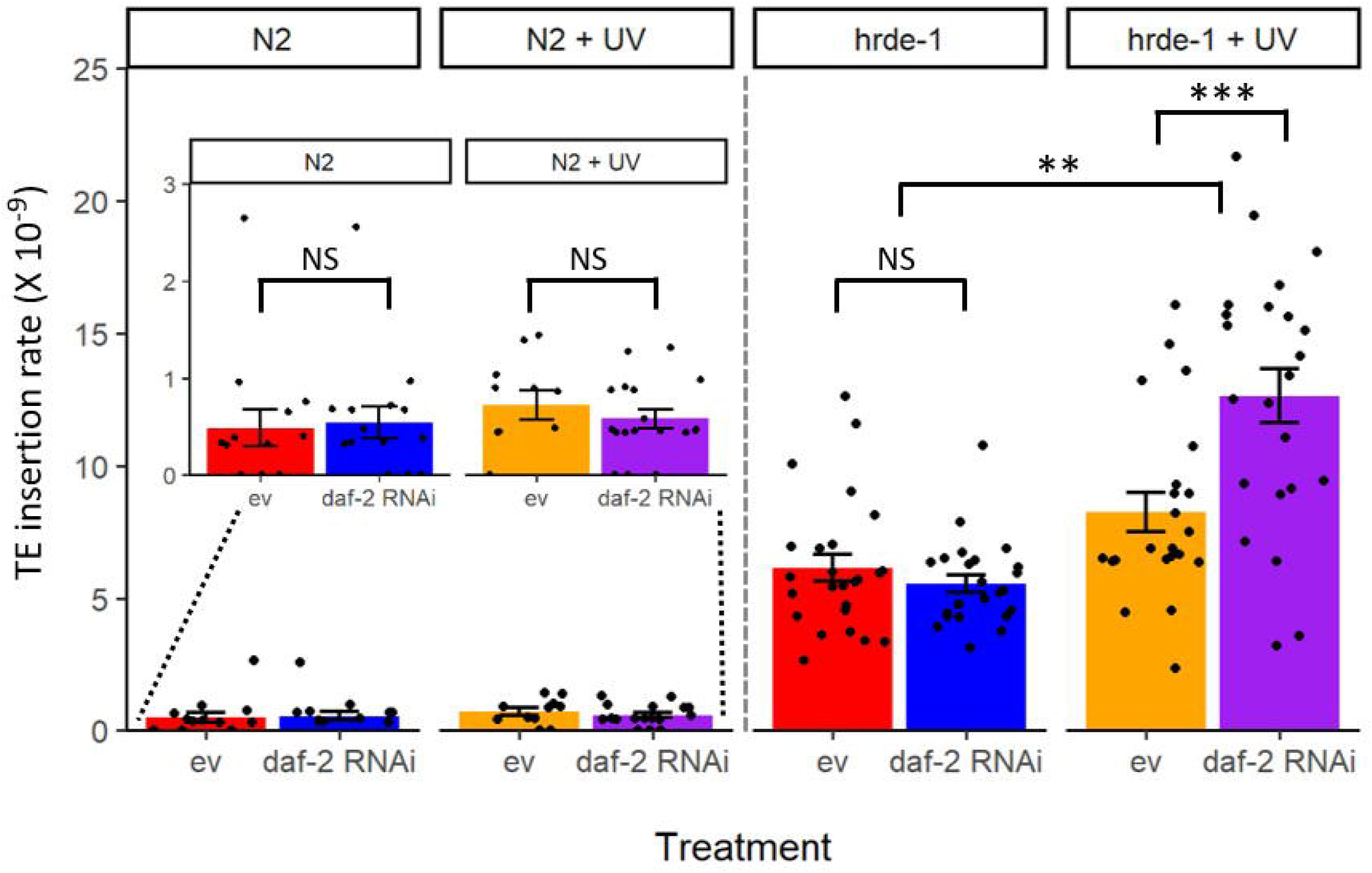
TE insertion rates of *C. elegans* mutation accumulation (MA) lines across two genetic backgrounds: N2 wild type and *hrde-1* mutant and four MA treatments: empty vector (ev) control (red), reduced insulin/IGF-1 signalling via adulthood *daf-2* RNAi (blue), ev + UV irradiation (orange) or *daf-2* RNAi + UV (purple). Bars represent mean ± standard error and individual points represent the line-specific TE insertion rates for individual MA lines, per MA generation, per callable genomic site. Statistical significance: NS, non-significant; * p<0.05; ** p<0.01; *** p<0.001.

While novel TE insertions were found in all *hrde-1* mutant MA lines, no TE insertions were found in 17 or 18% of UV-irradiated N2 MA lines and 27 or 36% of spontaneous N2 MA lines (for *daf-2* RNAi versus ev treatments, respectively). There was no difference in overall TE insertion rates between ev controls and rIIS among spontaneous N2 MA lines. However, fewer ev control lines had TE-driven insertions than rIIS lines, although the exact mechanisms are unclear.

Overall, the elevated germline TE-driven insertion rates in *hrde-1* mutants that are deficient in heritable RNAi-mediated silencing is in line with previous work that highlighted the role of Piwi piRNA silencing in determining germline TE activity (Sarkies et al., 2015; Pritam & Signor, 2025). Further increase in TE insertion rates under UV-induced MA in *hrde-1* mutants supports the UV-induced increase seen previously in *D. simulans* (Jardim et al., 2015), although we did not find a significant UV-induced increase in TE insertions under spontaneous MA. While one study in long-lived *D. melanogaster* IIS mutants found downregulation of fat body-specific TE expression under rIIS, the exact mechanisms were unclear (Tain et al., 2021) and we did not find a significant benefit of rIIS for reduced germline TE insertion rates.

### 2.4 Fine-scale spectrum of TE classes reveals effects of rIIS across genetic backgrounds

To determine whether there was an over-representation of certain TE classes when TE insertion rates were elevated in *hrde-1* mutants under rIIS and UV-induced MA, fine-scale spectra were examined. DNA transposons were the predominant class of TEs driving novel insertion mutations in all MA treatments, ranging from an average of 76 to 95% of total TE-driven insertion mutations (Figure 3). This is in support of previous work in *C. elegans* that suggested DNA transposons to be the most active (Sturm et al., 2023; Bessereau, 2016) and important contributor to spontaneous germline insertions (Moerman & Waterson, 1984; Eide & Anderson, 1985).

**Figure 3.**
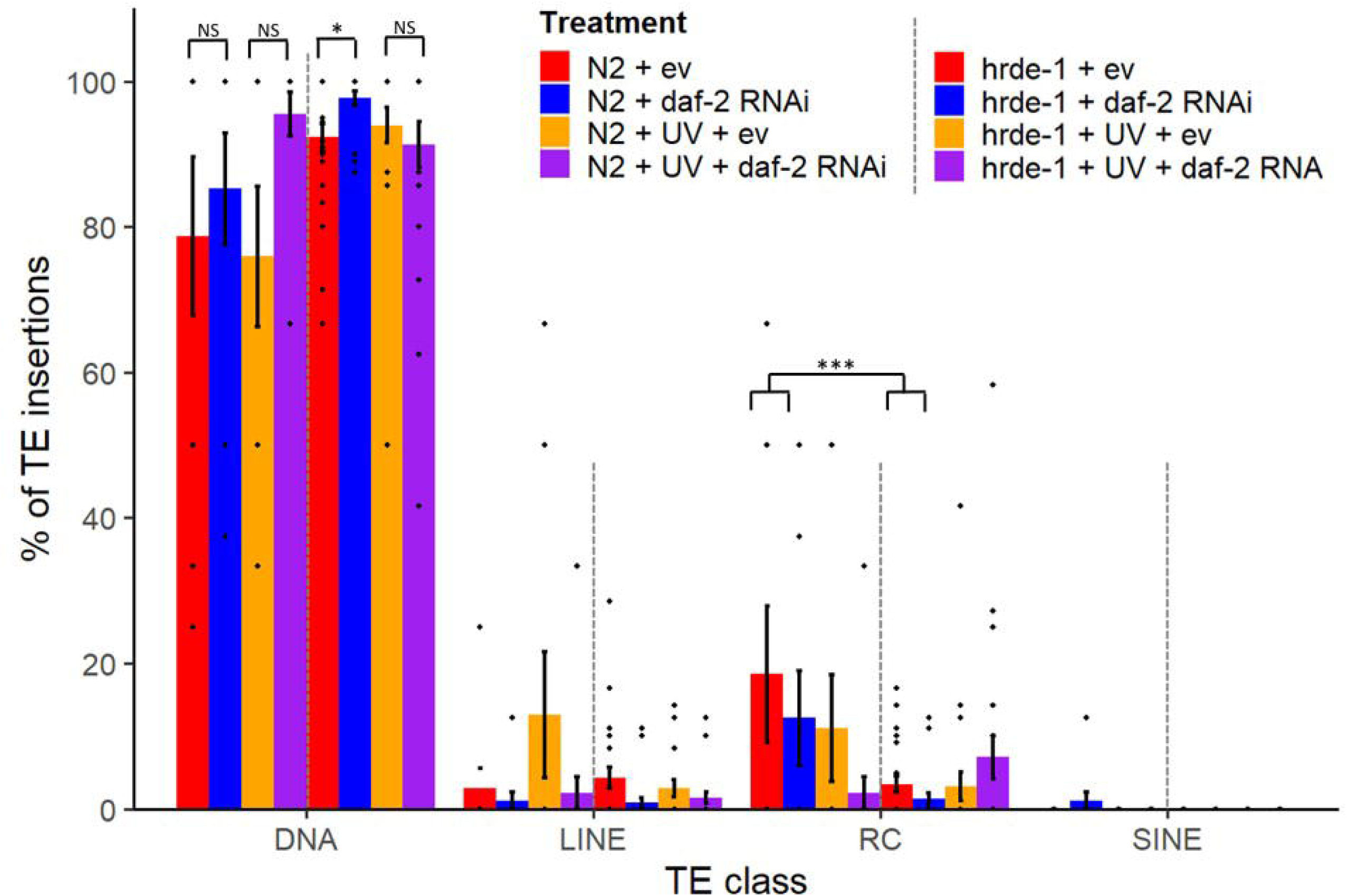
Percentage of novel TE insertions per MA line divided into classes (mean ± standard error). MA treatments indicate genetic background (N2 or *hrde-1* mutant, ‘hrde’), RNAi treatment (empty vector, ‘ev’ or *daf-2* RNAi in adulthood) and UV treatment (spontaneous MA or UV-induced MA). Individual points indicate the proportion of TEs in each class for individual MA lines with TE insertions. Statistical significance for key comparisons: NS, non-significant; * p<0.05; ** p<0.01; *** p<0.001.

Under rIIS, spontaneous *hrde-1* mutant MA lines had a greater proportion of insertion mutations driven by DNA transposons than for their respective ev controls (Binomial GLM, spontaneous *hrde-1* mutants: *z*=-2.224, *p*=0.026; Figure 3; Suppl. Table 8), even there was no significant difference in the TE insertion rates between these respective treatment (Figure 2). For *hrde-1* mutants under UV-induced MA, there was no significant effect of rIIS on the proportion of TE insertions that were DNA transposons (Binomial GLM, UV-irradiated *hrde-1* mutants: z=1.371, *p*=0.170; Figure 3), suggesting that the elevated TE insertion rates in this treatment (Figure 2) did not arise from an overrepresentation of DNA transposon-driven insertions. Curiously, the proportion of DNA transposon-driven insertions in UV-irradiated N2 MA lines on rIIS, was similar to the higher proportions of DNA transposons seen in *hrde-1* mutant MA lines, relative to the other N2 MA treatments (Figure 3; Supplementary Table 8). However, overall TE insertion rates in UV-irradiated N2 MA lines were still significantly lower than for the *hrde-1* mutant background (Figure 2).

TE insertions not classified as DNA transposons were either long interspersed element (LINE) retrotransposons, rolling circle (RC) transposons, or a very low proportion of short interspersed nuclear element (SINE) retrotransposons (Figure 3). Overall, few MA lines had novel non-DNA transposon-driven insertions (17% of lines had LINE, 22% of lines had RC and only one N2 spontaneous MA line had SINE-induced insertion mutations; Figure 3), which limited the statistical power to detect differences. N2 spontaneous MA lines had significantly greater proportions of RC-driven insertions than *hrde-1* mutant MA lines, with the exception of lower levels in N2 UV-irradiated MA lines under rIIS, that instead had a greater proportion of DNA transposon-driven insertions (Binomial GLM, Background *z*=5.777, *p*<0.001; Background x UV, *z*=-3.563, *p*<0.001; Figure 3; Suppl. Table 8). There was no significant difference detected in the proportion of LINE-driven TE insertions between MA treatments (Suppl. Table 8).

Overall, the higher TE insertion rates in *hrde-1* mutant MA lines were driven by a greater proportion of DNA transposon-driven insertions and a lower proportion of RC-driven insertions relative to N2 MA lines. While rIIS was associated with a greater proportion of DNA transposon-driven insertions in spontaneous *hrde-1* mutant MA lines and N2 UV-irradiated MA lines, there was no significant over-representation of a particular TE class in UV irradiated *hrde-1* mutant MA lines when TE insertion rates were elevated under rIIS.

### 2.5 Functional annotation of germline SNPs

We next asked whether rIIS changes the predicted functional impact of germline SNPs and whether such changes could help explain differences in MA line survival. We found no significant differences among MA treatments in the proportions of SNPs with high, moderate, low or modifier impact (Supplementary Table 9). Most SNPs (∼97%) fell into the modifier category, which includes non-coding variants with uncertain functional consequences. High, moderate and low impact SNPs accounted for 0.03 - 0.08%, 1.25 - 1.59% and 0.84 - 1.15% of mutations, respectively. Among SNPs with protein-coding effects, over 60% were missense, 30–40% synonymous and less than 3% nonsense. We found no significant effect of rIIS or UV irradiation on the proportions of missense or synonymous SNPs, and no significant effect of rIIS on the non-synonymous: synonymous ratio within pairwise treatment comparisons (Supplementary Figure 3, Supplementary Tables 10-11).

We then looked at functional enrichment among genes carrying these mutations. We found that genes carrying modifier SNPs commonly showed enrichment for development, metabolism and conserved longevity-regulating pathways, including mTOR. We also found enrichment for TGF-β, FoxO and Wnt signalling but this differed across treatments (please see Supplementary Figures 4-7). Interestingly, in spontaneous N2 MA lines, high and moderate impact SNPs under rIIS showed enrichment for biological regulation, mRNA surveillance and responses to metal ions, whereas controls showed enrichment for development and neuronal transport. On the other hand, under UV-induced MA, enrichment among high and moderate impact SNPs centred on cellular maintenance and biological regulation in rIIS lines, and neuronal function and cellular transport in controls. We found high impact variants in F31E3.6 and vab-19 under rIIS, F35A5.1 in controls, and *pals-26* and *tbc-17* in both treatments (Supplementary Figure 5).

In spontaneous *hrde-1* mutant MA lines, we found enrichment for biological regulation and protein processing among high and moderate impact SNPs under rIIS, and for cellular maintenance, protein modification, metabolism and dosage compensation in controls. We also found a high impact variant in F27C8.3 only under rIIS (Supplementary Figure 6).

Under UV-induced MA, high and moderate impact SNPs in *hrde-1* mutants on rIIS showed enrichment for development, reproduction, biological regulation, metabolism and cellular maintenance. Three high impact variants caused stop-codon loss in C33C12.1, *vab-19* and *sdha-1*, genes associated with neuronal function, embryonic and vulval development, and mitochondrial electron transport, respectively. Mutations in the human orthologue of sdha-1 are associated with Leigh syndrome. We found no significant functional enrichment among high and moderate impact SNPs unique to controls (Supplementary Figure 7).

These results identify possible genetic contributors to differences in MA line survival, including the rapid extinction of UV-irradiated *hrde-1* mutants under rIIS. Further detailed description of functional enrichment analysis is in the Supplementary Results.

### 2.6 Functional description of TE-driven insertions

To determine the potential impact of TE-driven insertions, we functionally described the genes into which they inserted, since there were insufficient genes for enrichment analyses. In non-irradiated N2 MA lines, TE insertions only overlapped with three genes for ev controls and two genes for MA lines on *daf-2* RNAi, encoding a membrane protein, a gustatory receptor and, two proteostasis-associated F-box proteins (T02G5.14 and Y46G5A.39), that have recently been linked with TE-driven regulatory changes (Almeida et al., 2025). In irradiated N2 MA lines on ev, TE insertions only overlapped with one gene, *srw-60*, encoding G protein-coupled receptor signalling pathway peptide associated with embryogenesis; but no TE insertions overlapped with genes for irradiated N2 MA lines on *daf-2* RNAi. Overall, since the number of genes with TE insertions was small for the N2 background, it was difficult to discern strong functional effects of rIIS.

For the *hrde-1* mutant MA lines, considerably more TE insertions overlapped with genes. Interestingly, for non-irradiated *hrde-1* mutants MA lines on ev, TE insertions occurred in genes associated regulation of the insulin signalling pathway (*daf-6*, *mpz-1*), methyltransferase activity (K01D12.1), mitochondrial unfolded protein response (*sphk-1*) and in a germline-expressed F-box domain gene (F08F3.6). In contrast, TE inserted into genes associated with nitrogen metabolism (*nit-1*) autophagy (*atg-4.1*) and neuronal function, in non-irradiated *hrde-1* mutants MA lines on *daf-2* RNAi. For irradiated lines on ev, TE insertions overlapped with genes involved in sensory perception (*srg-47*), encoding a sperm protein (Y105C5B.19), a prion-like germline-expressed gene (*pqn-20*) and a F-box domain gene (*fbxa-154*). TE insertions for irradiated *hrde-1* mutant MA lines on *daf-2* RNAi overlapped with genes associated with lipid metabolism (F46G10.4), protein folding (T22F3.12), and the negative regulation of apoptosis and cell cycle (*prk-1*). Overall, the greater range of important functions associated with TE insertions in the *hrde-1* mutant background, supports the previously reported role of *hrde-1* in promoting germline immortality (Buckley et al., 2012; Spracklin et al., 2017), and may have additionally contributed to their faster extinction under UV-induced MA, although the exact mechanisms underlying rIIS effects require further investigation.

### 2.7 Genomic location of germline SNPs

Mutation rates are expected to be lower in coding exonic sequences than in non-coding intronic sequences, due to selection for an increased efficiency of mismatch and transcription-coupled repair in exons, as observed in MA experiments (Krasovec et al. 2017; Konrad et al., 2019) and for human somatic mutation rates (Frigola et al., 2017). Here we asked whether rIIS reduces the proportion of germline mutations in coding regions, and whether this effect depends on functional HRDE-1. The null prediction was that mutations are expected to be distributed according to site mutability, chance and/or selection in controls. We hypothesised that rIIS may lead to lower GMRs via increased resource investment into germline repair of mutations and/or increased selection for the upregulation of gene expression in repair and resilience pathways that lower the resultant number and penetrance of germline mutations. This may involve alteration of the mutational process with respect to site mutability, selection and/or drift.

Across all MA treatments, SNPs were most frequently classed as upstream (35-38%) and downstream (38-40%) variants, with lower proportions of intron (12-15%), intergenic (6-7%), missense (1-2%) or synonymous (0.7-1%) variants (Suppl. Tables 12 & 13). Although SNPs occurred more frequently in introns than in intergenic regions or exons, their relative proportions differed between MA treatments (Suppl. Tables 12–14). Under rIIS (via *daf-2* RNAi), non-irradiated N2 MA lines had a significantly greater proportion of SNPs in introns relative to exons and intergenic regions, than for the respective ev control non-irradiated MA lines (Suppl. Table 14). This shift may reflect greater repair of coding regions and/or selection against deleterious coding mutations, potentially reducing the functional consequences of germline mutations. Conversely, non-irradiated *hrde-1* mutant MA lines on rIIS had significantly fewer SNPs occurring in introns than for respective ev control *hrde-1* mutant MA lines, relative to the proportion of SNPs occurring in exons for these MA treatments. For both N2 MA lines on the ev control and *hrde-1* mutant MA lines on *daf-2* RNAi, a greater proportion of SNPs occurred in introns relative to exons for UV-irradiated lines than for non-irradiated lines (Suppl. Table 14). Further detailed exploratory analyses of SNP genomic location are included in the Supplementary Results.

## 3 Conclusions

Our study demonstrates that reducing insulin signalling in adulthood significantly lowers GMR in *C. elegans*. While previous work found impaired germline stem cell proliferation from reduced IIS during development in *C.* elegans temperature-sensitive reduction-of-function *daf-2(e1370)* mutants (Michaelson et al., 2010; Cheng et al., 2024) and later in adult *daf-2(e1370)* mutants at 25°C (Narbonne et al., 2015), mediated in part via the somatic gonad (Cheng et al., 2024), the consequences of adulthood IIS knockdown for germline mutation load were unexplored. Direct measurement of the genomic impact of rIIS on the germline was important to better understand the underlying mechanisms, since *daf-2(e1370)* mutants also had reduced cell division and increased DAF-16/p53-dependent apoptosis in adulthood germline tumours (Pinkston et al., 2006), with improved oocyte quality (Luo et al., 2010) and delayed reproductive ageing in mated adults under rIIS (Hughes et al., 2007; Sultanova et al., 2025).

Because rIIS improves somatic maintenance and increases longevity, as well as increasing fitness of the surviving lines after 20 generations of UV-induced MA (Duxbury et al., 2022), our results challenge resource allocation theory, which predicts a trade-off between somatic maintenance and germline genome integrity (Maklakov & Immler, 2016). Our results suggest that insulin signalling regulates somatic and germline maintenance concordantly, and increased somatic maintenance under reduced IIS comes together with reduced germline mutation load. This result is broadly in line with the developmental theory of ageing which maintains that natural selection optimizes physiological processes for development, growth, and early-life reproduction but not for late-life function (Maklakov and Chapman 2019, Lemaitre et al 2024). One prediction of this theory is that selection optimizes gene expression for early-life function but not late-life function, and that gene expression during adulthood can optimised to slow down ageing and increase fitness (de Magalhaes and Church 2005, Maklakov and Chapman 2019, Lind et al. 2021, Lemaitre et al. 2024, Sultanova et al. 2025). The finding that somatic maintenance and germline mutation rate can be improved simultaneously by reducing insulin signalling in adulthood supports this view.

Earlier studies showed that relaxed selection results in the evolution of increased mutation rate in *C. elegans*, suggesting that directional selection to reduce mutation rate in *C. elegans* is opposed by genetic drift (Saxena et al., 2019). In small populations, the strength of drift exceeds selection, thus limiting the ability to purge deleterious mutations or fix alleles that are beneficial for fitness (Lynch, 2010; Sung et al., 2012). Our results suggest that a putative modifier allele that would reduce adult-only *daf-2* expression, mimicking our experimental setup, would reduce GMR without fitness costs. Essentially, this finding implies that either there is a hard insurmountable genetic constraint that prevents the evolution of age-specific *daf-2* expression, or that the benefits of reducing GMR in *C. elegans*, as found here, are insufficiently strong to overcome genetic drift in this system. The latter hypothesis is more parsimonious and is also in line with theoretical modelling, which suggests that genetic drift can maintain high mutation rates in selfing populations (Gervais & Roze, 2017).

The mechanism underlying the reduced GMR upon IIS downregulation implicates the nuclear Argonaute protein HRDE-1, suggesting an epigenetic pathway (Buckley et al., 2012) linking nutrient signalling directly to germline genomic stability. Previous work identified the role of functional HRDE-1 in the transgenerational silencing of transposable elements (Ashe et al., 2012), maintaining germline gametogenesis under high temperatures and promoting germline immortality (Buckley et al., 2012; Spracklin et al., 2017). Our work highlights a critical role for germline epigenetic regulation as a mediator between systemic physiological states, IIS and genome maintenance. This is likely to involve cross-talk between the IIS pathway and HRDE-1 function in the transgenerational silencing of germline TEs via piRNAs and small RNA inheritance. It is possible that IIS may also affect DNA methylation state or histone modification, with consequences for germline genome integrity and we encourage further work to elucidate the pathways involved. Consequently, our findings emphasize that epigenetic mechanisms play pivotal roles not only in developmental regulation but also in adaptive responses to environmental and metabolic conditions, shaping evolutionary trajectories (Heard & Martienssen, 2014).

Germline mutations in protein-coding genes were functionally enriched for development, reproduction, cellular maintenance and longevity-associated nutrient-sensing pathways (mTOR and IIS) in our *C. elegans* MA lines. We further identified several high impact inherited germline mutations in the UV irradiated MA lines with human orthologs that have been implicated in autoimmune diseases, renal cell carcinoma and rare neurodegenerative Leigh syndrome. These results reveal insights into the underlying mechanisms of MA line extinction and highlight the promising therapeutic value of strategies to reduce GMRs.

Collectively, our results (Duxbury et al., 2022, and here) refine and expand the evolutionary theory of ageing by emphasising the role of mutation-selection balance versus trade-off models (Hughes et al., 2002; Lemaitre et al., 2024). Specifically, these results suggest that age-specific gene expression can be modified to improve somatic maintenance, and late-life survival and reproduction and improve germline maintenance. Our findings underscore the evolutionary and translational potential of targeting IIS as a means to simultaneously enhance organismal longevity and reduce germline mutation load without compromising overall fitness, at least in some environments. Future research into the evolutionary conservation of this IIS-mediated genomic protection may open promising avenues for interventions aimed at reducing germline mutation burden in different organisms, potentially influencing strategies for mitigating age-related genomic deterioration.

## 4 Methods

### 4.1 Nematode strains and bacterial clones

The nematode (roundworm) *C. elegans* N2 wild-type (Bristol) and heritable RNAi deficiency 1 (*hrde-1)* mutant strains were defrosted from stocks acquired from Caenorhabditis Genetics Center (University of Minnesota, USA, funded by NIH Office of Research Infrastructure Programs, P40 OD010440) and from the lab of Prof. Eric Miska (University of Cambridge, UK), respectively, and stored at −80°C until use. The *hrde-1* mutant is derived from the N2 background.

To downregulate adulthood expression of the insulin-like sensing signaling receptor homolog, *daf-2*, we fed late-L4 larvae with *E. coli* bacteria expressing *daf-2* double-stranded RNA, which decreases mRNA levels of the complementary transcribed *daf-2* systemically (Fire et al., 1998; Timmons et al., 2001). The *daf 2* gene is upstream of and inhibits the action of master regulator daf-16/FOXO. RNase-III deficient, IPTG-inducible HT115 *E. coli* bacteria with an empty plasmid vector (L4440) was used as the control (as Timmons et al., 2001; Dillin et al., 2002; Duxbury et al., 2022). RNAi clones were acquired from the Vidal feeding library (Source BioScience, created by M. Vidal lab, Harvard Medical School, USA) and all clones were verified via sequencing prior to delivery.

### 4.2 Mutation accumulation lines

*Caenorhabditis elegans* mutation accumulation lines that we established in Duxbury et al. (2022), comprised of eight experimental treatments, that were the full factorial combination of RNAi (either reduced insulin/IGF-1 signalling via adulthood-only *daf-2* RNAi; or empty vector, ev), UV (either UV irradiated or non-irradiated, such that mutations accumulated spontaneously across generations) and genetic background (N2 wild type or *hrde-1* mutant). RNAi and UV treatments were applied each MA generation. There were two time-staggered experimental blocks consisting of all eight MA treatments. All MA lines were derived from respective N2 or *hrde-1* mutant founder populations.

Each MA line was propagated each generation using a single hermaphrodite offspring produced from the peak reproduction Day 2/Day 3 parent from the previous MA generation. By standardising parental age within each MA generation across all treatments, we controlled for variation in development or maturation time between experimental treatments. The single worm genetic bottlenecks each MA generation, allowed germline mutations to accumulate in the relative absence of selection, unless they led to extinction prior to reproduction. We avoided experimental selection on parental age across generations, due to alternation between Day 2 and Day 3 adult parents every other MA generation. More details about the mutation accumulation lines, their extinction trajectories across 40 generations of MA and their respective reproductive fitness after 20 generations of MA can be found in Duxbury et al. (2022).

### 4.3 Sample collection and DNA extractions

Nematode samples were collected from all MA line treatments: N2 + UV-irradiated (generation 25), N2 spontaneous MA (generation 35), *hrde-1* mutant spontaneous MA (generation 20) and *hrde-1* mutant UV-irradiated (generation 10 for ev and generation 6 for *daf-2* RNAi due to fast extinction), after the point of divergence of extinction trajectories for respective *daf-2* RNAi and ev treatments (Duxbury et al., 2022). The *daf-2* RNAi and empty vector samples were collected after the same number of generations of MA so that they could be directly compared. Each sample consisted of a flash-frozen pool of 20-40 adult Day 2-3 nematodes that were the F1 progeny of the respective focal MA line individual on Day 2 of adulthood. This avoided mutagenesis that can occur during population expansion for DNA extraction and subsequent bioinformatic filtering steps removed negligible somatic mutations that may have arisen. N2 and *hrde-1* mutant founder populations were also sequenced (three samples per founder population per genetic background). Sample sizes for the number of MA lines per treatment are listed in Table 1. Unequal sample sizes between treatments were due to insufficient DNA quantity or quality yielding from the DNA extractions for some samples, so they were excluded from whole genome sequencing.

Genomic DNA (gDNA) was extracted from each sample using an optimised protocol with the Qiagen Blood and Tissue Kit. Quantity and quality of extracted DNA was assessed with a Nanodrop spectrophotometer and Qubit quantification of double-stranded DNA; followed by gel electrophoresis. To obtain sufficient gDNA for sequencing we performed whole genome amplification with the Qiagen REPLI-g® Advanced DNA Single Cell Kit, that is optimised for <100ng of gDNA and highly uniform genome coverage. The kit uses multiple displacement amplification technology with an optimised, higher-fidelity Phi 29 polymerase formulation, which enables negligible sequence bias and minimal genomic dropouts. Product length was consistently greater than 10kb across samples.

### 4.4 Whole genome sequencing

Whole genome sequencing was performed with an Illumina NovaSeq 6000 S4 v1.5 flow cell with 150bp paired end reads, PCR-free technology (to reduce bias in allelic ratios) and bead-based size selection. Samples were split across two sequencing lanes, with 2,500 million reads per lane for each direction sequenced, to achieve a minimum of 50-60x sequencing depth (coverage) per sample.

### 4.5 Data processing

Raw reads in fastq format were individually adapter- and quality-trimmed using fastp/0.32.2 (Chen et al., 2018) and mapped against the latest version of the *C. elegans* N2 reference genome (WBcel235 NCBI, 2013, GenBank Accession no. GCA_000002985.3) using mem, BWA/0.7.17 (Li, 2013). The *C. elegans* genome is comprised of five autosomes, one sex chromosome and one mitochondrial chromosome, with total genome size of 100.3Mb.

Aligned reads were then sorted (using samtools/1.16.1 sort), indexed (using samtools/1.16.1 index; Danecek et al., 2021), and duplicates marked (using Picard/2.24.1 MarkDuplicates; http://broadinstitute.github.io/picard/). The resultant bam files were indexed (using samtools/1.16.1 index) and then used for mutation calling.

### 4.6 Variant calling

Single nucleotide polymorphisms (SNPs) were called relative to the relevant founder genetic background. To do this we first used “bcftools mpileup” and “bcftools call” to call variants in all MA lines relative to the *C. elegans* N2 reference genome (bcftools mpileup -a AD,ADF,ADR -r $chr -q 30 -Q 30 -f genome.fa -b $bam_list | bcftools call -mv -f GQ | bcftools filter -g 3 -G 3 -e ‘DP < 1100 || DP > 20000 || F_MISSING > 0.5 || QUAL < 30’ -Oz - o $output_file). We then created a consensus reference genome for each genetic background (N2 wild types and *hrde-1* mutants) by incorporating the SNPs called in the three founder lines per genetic background back into the *C. elegans* reference genome (using “bcftools consensus”). Finally, these newly-created consensus reference founder genomes were used to call variants in the MA lines relative to the relevant founder background. This ensured that variants called in the MA lines were not false positives or false negatives, but instead accounted for sites where the founder lines had changed since the *C. elegans* N2 reference genome.

SNPs were stringently filtered for quality (Supplementary Methods) and subsequently filtered to only include heterozygous SNPs. Heterozygous variants were genomic sites where there was one ancestral (founder) and one MA line-specific base substitution (SNP), relative to the founder. Since it was very unlikely that a homozygous SNP (base substitution in both copies at a specific genomic site) would occur after only 35 generations, it was most likely to be a false positive and filtered out (following Chen et al., 2023). Only SNPs that were unique to each MA line and not shared by any of the of MA lines in that UV and RNAi treatment combination, with both experimental blocks combined were included (using “bcftools isec”). These unique MA line-specific heterozygous SNPs, that were different both from the allele shared by the other lines and from the founder allele, were therefore assumed to be a line-specific germline mutation that occurred during the mutation accumulation process, thus further eliminating any false positives.

The “+split” plugin was used to create separate variant call files (VCFs) per MA line and “bcftools view” with the “--trim-alternate-alleles” command was used to produce VCFs containing only line-specific and heterozygous SNPs. VCFs were indexed using “bcftools index” prior to using “bcftools isec”. Indels and multiallelic variants (sites with more than one alternate allele) were filtered out from analyses, but insertion mutations that were induced by transposable elements were analysed separately (as below). Only variants of high quality and low stand bias were included (as above, and bcftools view -m 2 -M 2 --min-ac 3:minor -e ‘ALT=“*” || type!=“snp” || F_MISSING > 0.25’; bcftools +prune -m 0.6 -w20kb).

### 4.7 Germline mutation rates

Line-specific mutation rates (μ) were calculated as: μ = m/(L*n*T) (Denver et al., 2012; following Chen et al., 2023), where μ is the mutation rate per nucleotide site per generation for each MA line, m is the number of line-specific mutations (SNPs only), L is the number of MA lines (L = 1 for line-specific mutation rate), n is the number of nucleotide sites accessible for mutation calling (determined with “mosdepth v.0.3.2”, Pedersen & Quinlan 2018, from the .bam files by applying a coverage threshold between 13X and 118X inclusive, based on the mean coverage per sample of 39.45X; Supplementary Methods) and T is number of generations of MA. We excluded six N2 samples (three for *daf-2* RNAi and three for ev treatment) and one *hrde-1* mutant sample (from *daf-2* RNAi + UV treatment) that had less than 30% of the genome with callable sites, as these considerably skewed the mutation rates. On average, 85% of the genome had callable sites across the remaining samples. Data from the two experimental blocks was pooled due to uneven sample sizes between blocks and as their time-staggering (of a few days) was minor compared to the duration of 40 generations of mutation accumulation (over six months).

### 4.8 Mutational spectra

We determined the frequency of each type of base substitution using the outputs from “bcftools stats” for each MA line separately. Mutations were pooled into transitions (A:T- >G:C, G:C- >A:T) or transversions (A:T->C:G, G:C->T:A, A:T->T:A, G:C->C:G), and the transition:transversion ratios (Ts:Tv ratios) were calculated for each MA line.

### 4.9 Functional annotation

We used SnpEff (ver. 5.2.a; Cingolani et al., 2012a), SnpSift (Cingolani et al., 2012b) and ShinyGO (ver. 0.80; Ge et al., 2020), with annotation database WBcel235.105, using all protein-coding genes in the latest version of the *C. elegans* genome as the background, to perform functional enrichment analyses. We annotated SNPs on all six chromosomes (I-V, X) for all eight experimental treatments, but SnpEff was not able to annotate SNPs on the mitochondrial chromosome. This determined the impact of the SNPs (high, moderate, low or modifier), their functional class (nonsense, missense or silent), type (whether they occurred in exons, introns or upstream or downstream of genes), and the significantly enriched biological processes and KEGG pathways for SNP mutations (fold enrichment > 1.0, FDR < 0.05), with p-values corrected for multiple testing using the false discovery rate (FDR) computed using the Benjamini–Hochberg method. The 20 most enriched pathways were displayed (based on fold enrichment) to eliminate weakly enriched GO terms, thus controlling for potential false positives or biases. The null prediction was no difference in functional annotation between pairwise comparisons of MA treatments.

### 4.10 Transposable element (TE)-induced insertion mutations

TE-induced insertion mutations were identified from aligned reads in BAM files, separately for each MA line, using the Retroseq pipeline (Keane et al., 2013) and classified by TE type: DNA, LINE, SINE, rolling circle (RC) or long terminal repeat (LTR). TE insertions in MA lines that overlapped with those in the respective founder population were removed using “bedtools intersect” (option -v), to obtain only those that arose following the MA treatment. We then used “bedtools window” to select the subset of TE insertions that did not overlap within a 150bp window of a reference TE insertion, to identify novel TE insertions that occurred outside of the proximity of a previously identified TEs. Window size was set as the insert size of the sequencing libraries (as recommended by Keane et al., 2013). Finally, after comparing different quality-filtering thresholds (Supplementary Methods) and finding qualitatively similar patterns of TE numbers between treatments, we used stringent quality filtering on novel TEs (recommended number of supporting reads, GQ>20; maximum breakpoint criteria, FL=8; Keane et al., 2013), to obtain only the highest confidence TEs. MA line-specific TE insertion rates were calculated as the number of TE insertions per MA line, per generation, relative to the number of callable genomic sites (as for line-specific SNP mutation rates). We used WormBase to provide functional descriptions of the small number of TEs that inserted into genes.

### 4.11 Statistical analyses

All statistical analyses were performed in R (ver. 4.3.1; R Core Team, 2023). Mutation rates (for SNPs and for TE insertions separately) were first analysed using a combined generalised linear model with Gaussian error structure, with RNAi treatment, UV treatment, genetic background (either N2 or *hrde-1* mutant) and their three-way and two-way interactions, fitted as explanatory factors. Fitting experimental block as a random effect in a Gaussian linear mixed effects model (lmer) did not qualitatively change the results, but provided reduced model fit (using AICtab) so data from both experimental blocks was combined for all analyses (without random block effects).

Secondly, pairwise comparisons were made between the number of mutations in *daf-2* RNAi versus empty vector treatments, after the data had been subset first by genetic background and then by UV treatment. For example, the number of mutations in N2 UV irradiated MA lines were compared between *daf-2* RNAi versus empty vector treatments, such that there were an identical number of generations of MA within each pairwise comparison (as Table 1). Only for irradiated *hrde-1* mutants was it not possible to match the number of MA generations (10 generations for empty vector, 6 generations for *daf-2* RNAi, due to the rapid extinction of the latter) so these were omitted from the analysis of number of mutations. To analyse the number of mutations we fitted seven different generalised linear models: Poisson (with or without an offset for the log-transformed number of callable genomic sites); then to account for overdispersion, either quasipoisson, or negative binomial (with or without an offset for the log-transformed number of callable loci), and finally Binomial or Quasibinomial (with the number of mutations divided by the number of callable sites as the response variable and the number of callable sites also added as a weight). Using “AICtab” in the bbmle package we found that the negative binomial model with offset had best fit for all pairwise comparisons.

We determined whether the spectra of point mutations differed from the expected Ts/Tv ratio of 0.5 using a one-sided, paired Wilcoxon signed rank test, as differences between actual and expected ratios did not satisfy the normality assumption. Mutational spectra were compared between the experimental treatments using a Gaussian GLM and fine-scale mutational spectra with a GLM with quasibinomial error structure to account for overdispersion (best fit with AICtab). Pairwise comparisons of the functional impact of protein-coding SNPs between RNAi treatments, and of the proportions of SNPs occurring in exons, introns or intergenic regions, were analysed with a two-tailed z-test with Yates continuity correction (prop.test in R).

## Acknowledgements

We thank Dr.s Hwei-yen Chen (Lund University, Sweden), Martyna Zwoińska (Uppsala University), Daniel Marcu (UEA) and Adam Ciezarek (Earlham Institute, Norwich) for helpful advice on the bioinformatics analyses. We also thank Dr.s Hwei-yen Chen and Cecile Jolly (Uppsala University, Sweden) for advice on the nematode gDNA extractions, and Dr. Kris Sales (UEA) for assistance with nematode collection. Finally, we thank Dr. Zahida Sultanova (UEA) for sharing gene expression data for chromosomal location overlap analyses prior to publication. N2 nematode strains were provided by the CGC, which is funded by NIH Office of Research Infrastructure Programs (P40 OD010440). We thank Wormbase for providing access to the *C. elegans* genome, annotation and genetic resources. The research presented in this paper was carried out on the High Performance Computing Cluster supported by the Research and Specialist Computing Support service at the University of East Anglia.

## Author Contributions

Conceptualization: E.M.L.D, A.A.M., S.I and H.C., methodology: E.M.L.D, A.A.M, S.I., H.C., A.M.G and J.-C. D. C., data analysis: E.M.L.D. and A.M.G., writing – original draft: E.M.L.D. and A.A.M., writing – review and editing: E.M.L.D., A.A.M., A.M.G., S.I., J.-C. D. C. and H.C.

## Data and code availability

All code for bioinformatics and statistical analyses are available in GitHub (link shared with reviewers). Data from bioinformatics pipelines is available on FigShare (private link for reviewers; will be freely accessible upon publication of the manuscript). Raw sequence data files are available on the European Nucleotide Archive (Project: PRJEB102952, currently private embargo, public link upon publication).

## References

Agrawal, A.F., and Wang, A.D. (2008). Increased transmission of mutations by low-condition females: Evidence for condition-dependent DNA repair. PLoS Biol. 6, 389–395.

Almeida MV, Li Z, Rebelo-Guiomar P, Dallaire A, Fiedler L, Price JL, Sluka J, Liu X, Butter F, Rödelsperger C, Miska EA. Transposable Elements Drive Regulatory and Functional Innovation of F-box Genes. Mol Biol Evol. 2025;42(5):msaf097.

Antebi A. Regulation of longevity by the reproductive system. Exp Gerontol. 2013; 48:596–602.

Arantes-Oliveira, N., Apfield, J., Dillin, A. & Kenyon, C. 2002. Regulation of life-span by germ-line stem cells in Caenorhabditis elegans. Science, 295, 502–505.

Ashe, A.; Sapetschnig, A.; Weick, E.-M.; Mitchell, J.; Bagijn, M.P.; Cording, A.C.; Doebley, A.-L.; Goldstein, L.D.; Lehrbach, N.J.; Le Pen, J.;, et al. piRNAs Can Trigger a Multigenerational Epigenetic Memory in the Germline of C. elegans. Cell 2012, 150, 88–99.

Babbitt GA, Cotter CR. 2011. Functional conservation of nucleosome formation selectively biases presumably neutral molecular variation in yeast genomes. Genome Biol Evol 3: 15–22. 10.1093/gbe/evq081

Berger D, Stangberg J, Grieshop K, Martinossi-Allibert I, Arnqvist G. 2017 Temperature effects on life-history trade-offs, germline maintenance and mutation rate under simulated climate warming. Proc. R. Soc. B 284: 20171721.

Bessereau, J.-L. Transposons in C. elegans, WormBook (The C. elegans Research Community, WormBook, 2016).

Biglou SG, Bendena WG, Chin-Sang I. An overview of the insulin signaling pathway in model organisms Drosophila melanogaster and Caenorhabditis elegans. Peptides. 2021;145:170640.

Buckley, B.A., Burkhart, K.B., Gu, S.G., Spracklin, G., Kershner, A., Fritz, H., Kimble, J., Fire, A. & Kennedy, S.. 2012. A nuclear Argonaute promotes multigenerational epigenetic inheritance and germline immortality. Nature, 489, 447–451.

Chen S, Zhou Y, Chen Y, Gu J. fastp: an ultra-fast all-in-one FASTQ preprocessor. Bioinformatics. 2018;34(17):i884–i890.

Chen HY, Jolly C, Bublys K, Marcu D, Immler S. Trade-off between somatic and germline repair in a vertebrate supports the expensive germ line hypothesis. Proc Natl Acad Sci U S A. 2020;117(16):8973–8979.

Chen HY, Krieg T, Mautz B, Jolly C, Scofield D, Maklakov AA, Immler S. Germline mutation rate is elevated in young and old parents in *Caenorhabditis remanei*. Evol Lett. 2023;7(6):478–489.

Cheng E, Lu R, Gerhold AR. Non-autonomous insulin signaling delays mitotic progression in C. elegans germline stem and progenitor cells. PLoS Genet. 2024 Dec 23;20(12):e1011351. doi: 10.1371/journal.pgen.1011351.

Cingolani P, Platts A, Wang le L, Coon M, Nguyen T, Wang L, Land SJ, Lu X, Ruden DM. 2012a A program for annotating and predicting the effects of single nucleotide polymorphisms, SnpEff: SNPs in the genome of Drosophila melanogaster strain w1118; iso-2; iso-3. Fly (Austin);6(2):80–92.

Cingolani P, Patel VM, Coon M, Nguyen T, Land SJ, Ruden DM, Lu X. 2012b Using Drosophila melanogaster as a Model for Genotoxic Chemical Mutational Studies with a New Program, SnpSift. Front Genet. 15; 3–35.

Danecek P, Bonfield JK, Liddle J, Marshall J, Ohan V, Pollard MO, Whitwham A, Keane T, McCarthy SA, Davies RM, Li H. Twelve years of SAMtools and BCFtools. Gigascience. 2021;10(2):giab008.

de Magalhaes, J.P. & Church, G.M. 2005. Genomes optimize reproduction: aging as a consequence of the developmental program. Physiology, 20, 252–259.

Dennis S, Sheth U, Feldman JL, English KA, Priess JR. 2012. C. elegans germ cells show temperature and age dependent expression of Cer1, a gypsy/Ty3-related retrotransposon. PLOS Pathogens 8:e1

Denver DR, Dolan PC, Wilhelm LJ, Sung W, Lucas-Lledo JI, Howe DK, Lewis SC, Okamoto K, Thomas WK, Lynch M, et al. 2009. A genome-wide view of *Caenorhabditis elegans* base-substitution mutation processes. Proc Natl Acad Sci USA 106: 16310–16314. 10.1073/pnas.0904895106

Denver, D. R., L. J. Wilhelm, D. K. Howe, K. Gafner, P. C. Dolan et al., 2012 Variation in base-substitution mutation in experimental and natural lineages of Caenorhabditis nematodes. Genome Biol. Evol. 4: 513–522.

Dillin A, Crawford DK, Kenyon C. Timing requirement for insulin/IGF-1 signalling in C elegans. Science. 2002;298:830–4.

Duxbury EML, Carlsson H, Sales K, Sultanova Z, Immler S, et al. Multigenerational downregulation of insulin/ IGF-1 signalling in adulthood improves lineage survival, reproduction, and fitness in Caenorhabditis elegans supporting the developmental theory of ageing. Evolution. 2022;76:2829–45.

Duxbury EML, Carlsson H, Kimberley A, Ridge Y, Johnson K, Maklakov AA. Reduced insulin/IGF-1 signalling upregulates two anti-viral immune pathways, decreases viral load and increases survival under viral infection in C. elegans. Geroscience. 2024;46(6):5767–5780.

Eide D, Anderson P (1985) The gene structures of spontaneous mutations affecting a Caenorhabditis elegans myosin heavy chain gene. Genetics 109: 67–79.

Ermolaeva, M.A., Segref, A., Dakhovnik, A., Ou, H.-L., Schneider, J.I., Utermoehlen, O., Hoppe, T., and Schumacher, B. (2013). DNA damage in germ cells induces an innate immune response that triggers systemic stress resistance. Nature 501, 416–420.

Evans EA, Chen WC, Tan M-W. The DAF-2 insulin-like signaling pathway independently regulates aging and immunity in C elegans. Aging Cell. 2008;7:879–93.

Fire, A., Xu, S., Montgomery, M.K., Kostas, S.A., Driver, S.E. & Mello, C.C.. 1998. Potent and specific genetic interference by double-stranded RNA in Caenorhabditis elegans. Nature, 391, 806–811.

Frigola J, Sabarinathan R, Mularoni L, Muiños F, Gonzalez-Perez A, López-Bigas N. Reduced mutation rate in exons due to differential mismatch repair. Nat Genet. 2017; 49(12):1684–1692. doi: 10.1038/ng.3991.

Furió, V., Moya, A., & Sanjuán, R. (2005). The cost of replication fidelity in an RNA virus. Proceedings of the National Academy of Sciences, 102, 10233–10237.

Garsin DA, Villanueva JM, Begun J, Kim DH, Sifri CD, et al. Long-lived C. elegans daf-2 mutants are resistant to bacterial pathogens. Science. 2003;300:1921.

Ge SX, Jung D, Yao R. ShinyGO: a graphical gene-set enrichment tool for animals and plants. Bioinformatics. 2020;36(8):2628–2629.

Gervais C, Roze D. Mutation Rate Evolution in Partially Selfing and Partially Asexual Organisms. Genetics. 2017 Dec;207(4):1561–1575.

Guo C, McDowell IC, Nodzenski M, Scholtens DM, Allen AS, Lowe WL, Reddy TE. 2017. Transversions have larger regulatory effects than transitions. BMC Genomics 18: 394. 10.1186/s12864-017-3785-4

Heard E, Martienssen RA. Transgenerational epigenetic inheritance: myths and mechanisms. Cell. 2014;157:95–109.

Holzenberger M, Dupont J, Ducos B, Leneuve P, Geloen A, Even PC, et al. IGF-1 receptor regulates lifespan and resistance to oxidative stress in mice. Nature. 2003;421(6919):182–7.

Honda Y, Honda S. The daf-2 gene network for longevity regulates oxidative stress resistance and Mn-superoxide dismutase gene expression in Caenorhabditis elegans. FASEB J. 1999;13(11):1385–93.

Hsin, H. & Kenyon, C. 1999. Signals from the reproductive system regulate the lifespan of C. elegans. Nature, 399, 362–366.

Hughes KA, J.A. Alipaz, J.M. Drnevich, & R.M. Reynolds, A test of evolutionary theories of aging, Proc. Natl. Acad. Sci. U.S.A., 2002; 99 (22) 14286–14291.

Hughes SE, Evason K, Xiong C, Kornfeld K. Genetic and pharmacological factors that influence reproductive aging in nematodes. PLoS Genet. 2007 Feb 16;3(2):e25. doi: 10.1371/journal.pgen.0030025

Jardim, S. S., Schuch, A. P., Pereira, C. M. & Loreto, E. L. S. Effects of heat and UV radiation on the mobilization of transposon mariner Mos1. Cell Stress Chaperones 20, 843–851 (2015).

Keane TM, Wong K, Adams DJ. RetroSeq: transposable element discovery from next-generation sequencing data. Bioinformatics. 2013;29(3):389–90.

Kenyon, C. The genetics of ageing. Nature 464, 504–512 (2010).

Kimura, M. (1967). On the evolutionary adjustment of spontaneous mutation rates. Genetics Research, 9(1), 23–34.

Konrad A, Brady MJ, Bergthorsson U, Katju V. Mutational Landscape of Spontaneous Base Substitutions and Small Indels in Experimental *Caenorhabditis elegans* Populations of Differing Size. Genetics. 2019;212(3):837–854.

Krasovec M, Eyre-Walker A, Sanchez-Ferandin S, Piganeau G. Spontaneous Mutation Rate in the Smallest Photosynthetic Eukaryotes. Mol Biol Evol. 2017 Jul 1;34(7):1770–1779. doi: 10.1093/molbev/msx119. PMID: 28379581; PMCID: PMC5455958.

Lemaitre J-F, Moorad J, Gaillard J-M, Maklakov AA, Nussey DH (2024) A unified framework for evolutionary genetic and physiological theories of aging. PLoS Biol 22(2): e3002513.

Li H. (2013) Aligning sequence reads, clone sequences and assembly contigs with BWA-MEM. arXiv:1303.3997v2

Lind MI, Ravindran S, Sekajova Z, Carlsson H, Hinas A, et al. Experimentally reduced insulin/IGF-1 signalling in adulthood extends lifespan of parents and improves Darwinian fitness of their offspring. Evol Lett. 2019;4:737–44.

Lind, M. I., Carlsson, H., Duxbury, E. M. L., Ivimey-Cook, E., & Maklakov, A. A. (2021). Cost-free lifespan extension via optimization of gene expression in adulthood aligns with the developmental theory of ageing. Proceedings of the Royal Society B: Biological Sciences, 288, 20201728.

Lind MI, Mautz BS, Carlsson H, Hinas A, Gudmunds E, Maklakov AA. Sex-specific growth and lifespan effects of germline removal in the dioecious nematode Caenorhabditis remanei. Aging Cell. 2024;23(11):e14290.

Lithgow GJ, White TM, Melov S, Johnson TE. Thermotolerance and extended life-span conferred by single-gene mutations and induced by thermal stress. Proc Natl Acad Sci U S A. 1995;92(16):7540–4.

Luo S, Kleemann GA, Ashraf JM, Shaw WM, Murphy CT. TGF-β and insulin signaling regulate reproductive aging via oocyte and germline quality maintenance. Cell. 2010 Oct 15;143(2):299–312. doi: 10.1016/j.cell.2010.09.013

Lynch M. 2010 Evolution of the mutation rate. Trends Genet. 26, 345–352.

Maklakov AA, Immler S. The Expensive Germline and the Evolution of Ageing. Curr Biol. 2016; 26(13):R577–R586.

Maklakov, A.A. & Chapman, T. 2019. Evolution of ageing as a tangle of trade-offs: energy versus function. Proc. R. Soc. B Biol. Sci., 286, 20191604.

Meier B, Cooke SL, Weiss J, Bailly AP, Alexandrov LB, Marshall J, Raine K, Maddison M, Anderson E, Stratton MR, Gartner A, Campbell PJ. C. elegans whole-genome sequencing reveals mutational signatures related to carcinogens and DNA repair deficiency. Genome Res. 2014;24(10):1624–36.

Meier B, Volkova NV, Hong Y, Schofield P, Campbell PJ, Gerstung M, Gartner A. Mutational signatures of DNA mismatch repair deficiency in *C. elegans* and human cancers. Genome Res. 2018;28(5):666–675.

Michaelson D, Korta DZ, Capua Y, Hubbard EJ. Insulin signaling promotes germline proliferation in C. elegans. Development. 2010;137:671–80.

Moerman DG, Waterston RH (1984) Spontaneous unstable unc-22 IV mutations in C. elegans var. Bergerac. Genetics 108: 859–877.

Moses, E., Atlan, T., Sun, X., Franěk, R., Siddiqui, A., Marinov, G. K., Shifman, S., Zucker, D. M., Oron-Gottesman, A., Greenleaf, W. J., Cohen, E., Ram, O., & Harel, I. (2024). The killifish germline regulates longevity and somatic repair in a sex-specific manner. Nat. Aging, 4, 1–23.

Murphy CT, Hu PJ. Insulin/insulin-like growth factor signaling in C. elegans. WormBook. 2013 Dec 26:1–43. doi: 10.1895/wormbook.1.164.1.

Narbonne P, Maddox PS, Labbé JC. DAF-18/PTEN locally antagonizes insulin signalling to couple germline stem cell proliferation to oocyte needs in C. elegans. Development. 2015 Dec 15;142(24):4230–41. doi: 10.1242/dev.130252.

Pedersen BS, Quinlan AR. Mosdepth: quick coverage calculation for genomes and exomes. Bioinformatics. 2018;34(5):867–868.

Pinkston JM, Garigan D, Hansen M, Kenyon C. Mutations that increase the life span of C. elegans inhibit tumor growth. Science. 2006 Aug 18;313(5789):971–5. doi: 10.1126/science.1121908.

Poetsch AR, Boulton SJ, Luscombe NM. 2018. Genomic landscape of oxidative DNA damage and repair reveals regioselective protection from mutagenesis. Genome Biol 19: 215. 10.1186/s13059-018-1582-2

Pritam S., Signor S. Evolution of piRNA-guided defense against transposable elements. Trends in Genetics 2025; 41(5): P390–401.

R Core Team (2023) R: A Language and Environment for Statistical Computing. R Foundation for Statistical Computing, Vienna, Austria. https://www.R-project.org/

Rajaei M, Saxena AS, Johnson LM, Snyder MC, Crombie TA, Tanny RE, Andersen EC, Joyner-Matos J, Baer CF. Mutability of mononucleotide repeats, not oxidative stress, explains the discrepancy between laboratory-accumulated mutations and the natural allele-frequency spectrum in *C. elegans*. Genome Res. 2021;31(9):1602–1613.

Regan JC, Froy H, Walling CA, Moatt JP, Nussey DH. Dietary restriction and insulin-like signalling pathways as adaptive plasticity: A synthesis and re-evaluation. Funct Ecol. 2020; 34:107–128.

Sarkies P, Selkirk ME, Jones JT, Blok V, Boothby T, Goldstein B, et al. (2015) Ancient and Novel Small RNA Pathways Compensate for the Loss of piRNAs in Multiple Independent Nematode Lineages. PLoS Biol 13(2): e1002061.

Saxena AS, Matthew P Salomon, Chikako Matsuba, Shu-Dan Yeh, Charles F Baer, Evolution of the Mutational Process under Relaxed Selection in *Caenorhabditis elegans*, Molecular Biology and Evolution, 2019; 36: 239–251.

Shemesh, N., Shai, N. & Ben-Zvi, A. 2013. Germline stem cell arrest inhibits the collapse of somatic proteostasis early in Caenorhabditis elegans adulthood. Aging Cell, 12, 814–822.

Sniegowski PD, Gerrish PJ, Johnson T, Shaver A. The evolution of mutation rates: separating causes from consequences. Bioessays. 2000;22:1057–66.

Spracklin, G., Fields, B., Wan, G., Becker, D., Wallig, A., Shukla, A. & Kennedy, S. 2017. The RNAi inheritance machinery of *Caenorhabditis elegans*. Genetics, 206, 1403–1416.

Sturm Á, Saskői É, Hotzi B, Tarnóci A, Barna J, Bodnár F, Sharma H, Kovács T, Ari E, Weinhardt N, Kerepesi C, Perczel A, Ivics Z, Vellai T. Downregulation of transposable elements extends lifespan in Caenorhabditis elegans. Nat Commun. 2023;14(1):5278.

Sultanova Z, Shen A, Hencel K, Carlsson H, Crighton Z, Clifton D, Akay A, Maklakov AA. Optimising Age-Specific Insulin Signalling to Slow Down Reproductive Ageing Increases Fitness in Different Nutritional Environments. Aging Cell. 2025:e14481.

Sun, L. et al. Heat stress-induced transposon activation correlates with 3D chromatin organization rearrangement in Arabidopsis. Nat. Commun. 11, 1886 (2020).

Sung W, Ackerman MS, Miller SF, Doak TG, Lynch M. 2012. Drift-barrier hypothesis and mutation rate evolution. PNAS 109(45):18488–92.

Suzuki T, Kamiya H. 2017. Mutations induced by 8-hydroxyguanine (8-oxo-7,8-dihydroguanine), a representative oxidized base, in mammalian cells. Genes Environ 39: 2. 10.1186/s41021-016-0051-y

Tain LS, Sehlke R, Meilenbrock RL, Leech T, Paulitz J, Chokkalingam M, Nagaraj N, Grönke S, Fröhlich J, Atanassov I, Mann M, Beyer A, Partridge L. Tissue-specific modulation of gene expression in response to lowered insulin signalling in *Drosophila*. Elife. 2021 Apr 21;10:e67275.

Thondamal, M., Witting, M., Schmitt-Kopplin, P. & Aguilaniu, H. 2014. Steroid hormone signalling links reproduction to lifespan in dietary restricted Caenorhabditis elegans. Nat. Commun., 5, 4879.

Timmons, L., Court, D.L. & Fire, A.. 2001. Ingestion of bacterially expressed dsRNAs can produce specific and potent genetic interference in Caenorhabditis elegans. Gene, 263, 103–112.

