## Supplementary Materials for "Lifespan-extending downregulation of insulin signalling reduces germline mutation load"

### Supplementary Material

#### Supplementary Figures

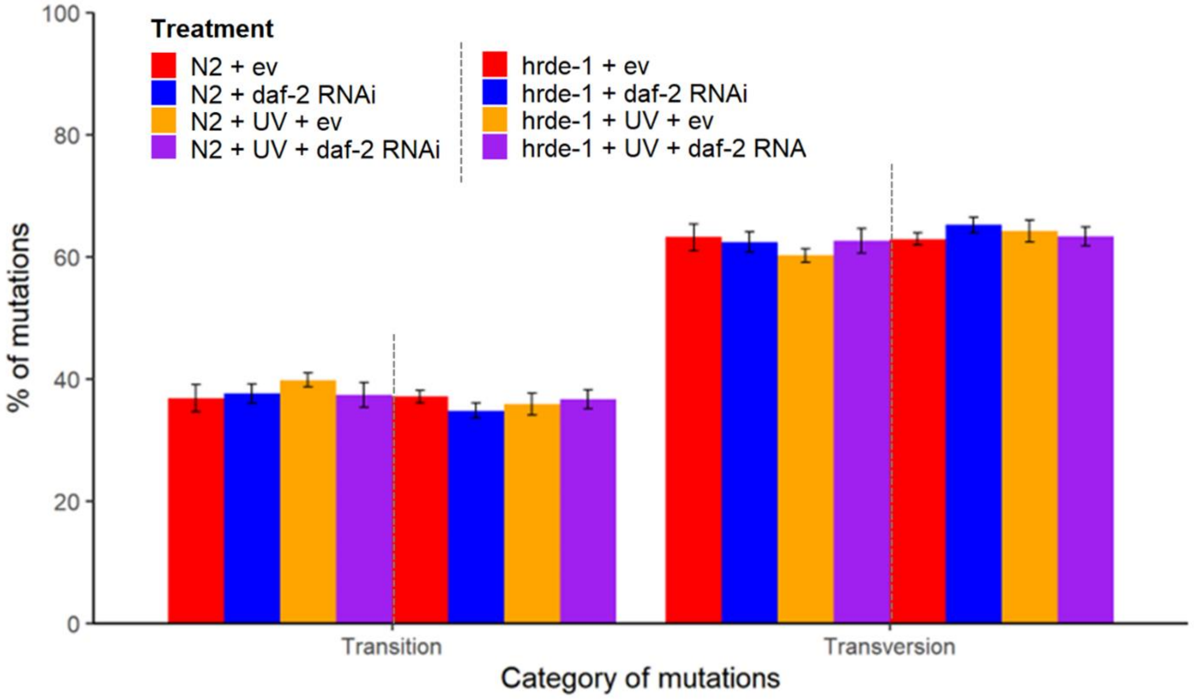


**Supplementary Figure 1. Percentage of mutations per MA treatment divided into transitions or transversions (mean ± standard error).** MA treatments indicate genetic background (N2 wild type or *hrde-1* mutant, ‘hrde-1’), RNAi treatment (empty vector, ‘ev’ or *daf-2* RNAi in adulthood) and UV treatment (spontaneous MA or UV-induced MA).

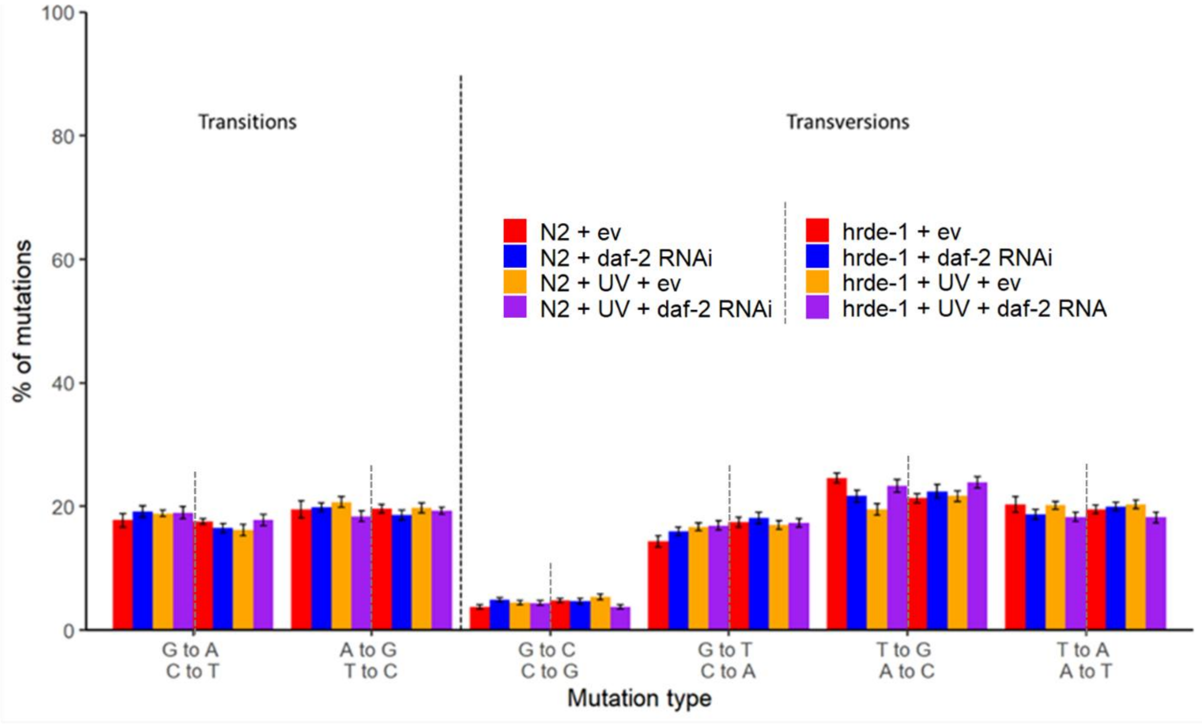


**Supplementary Figure 2. Percentage of mutations per treatment divided into point mutation type for transitions and transversions (mean ± 1 s.e.) per MA treatment.** MA treatments indicate genetic background (N2 or *hrde-1* mutant, 'hrde'), RNAi treatment (empty vector, 'ev' or *daf-2* RNAi in adulthood) and UV treatment (spontaneous MA or UV-induced MA).

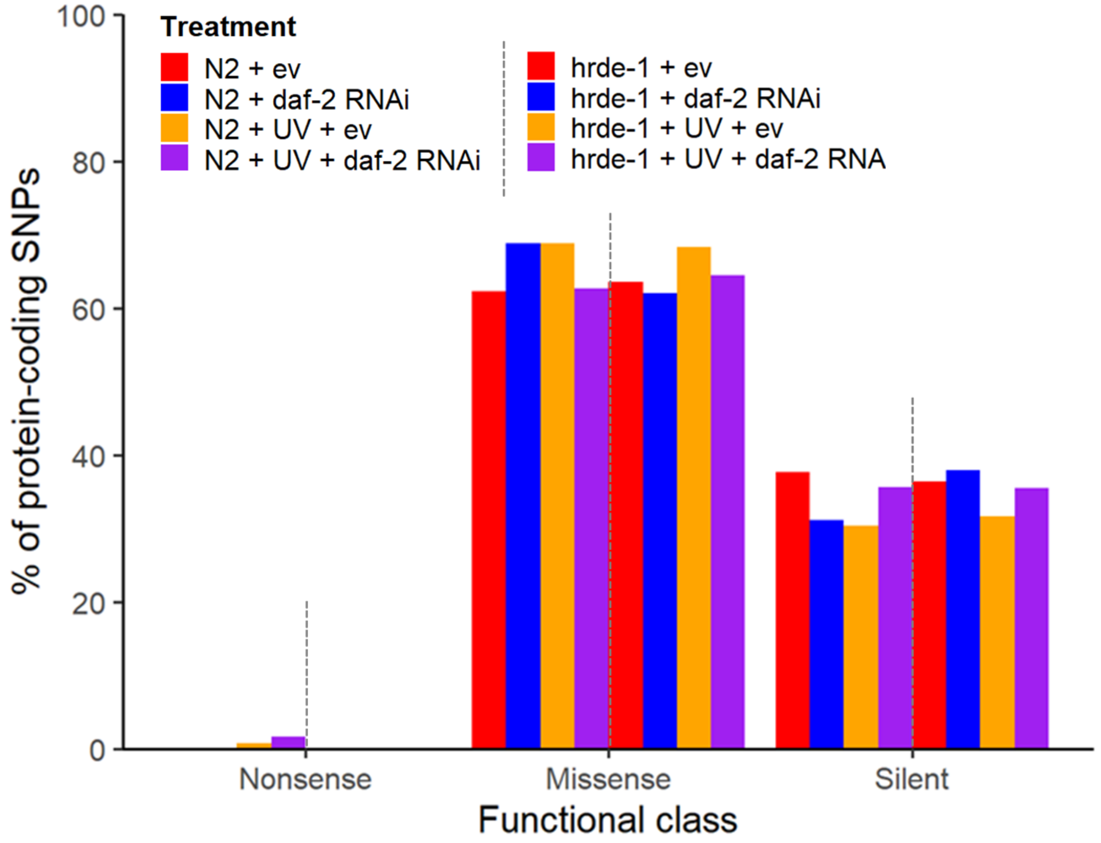


**Supplementary Figure 3. Functional class of protein-coding single nucleotide polymorphism (SNP) mutations per MA treatment.** MA treatments indicate genetic background (N2 or *hrde-1* mutant, ‘hrde’), RNAi treatment (empty vector, ‘ev’ or *daf-2* RNAi in adulthood) and UV treatment (spontaneous MA or UV-induced MA).

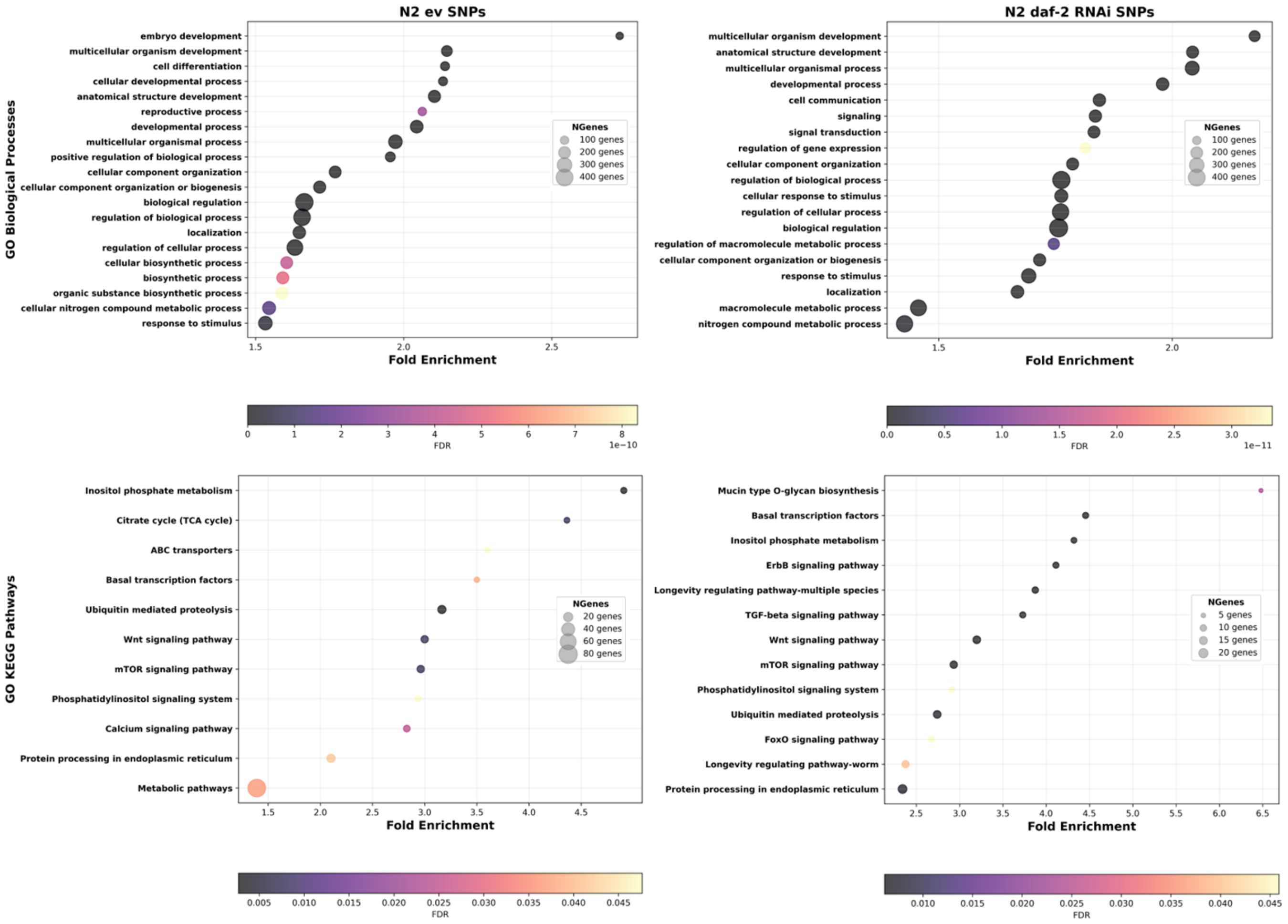


**Supplementary Figure 4.** Functional enrichment analysis of all single nucleotide polymorphism (SNP) mutations in N2 wild type spontaneous MA lines under reduced adulthood insulin signalling, via *daf-2* RNAi and in empty vector (ev) controls.

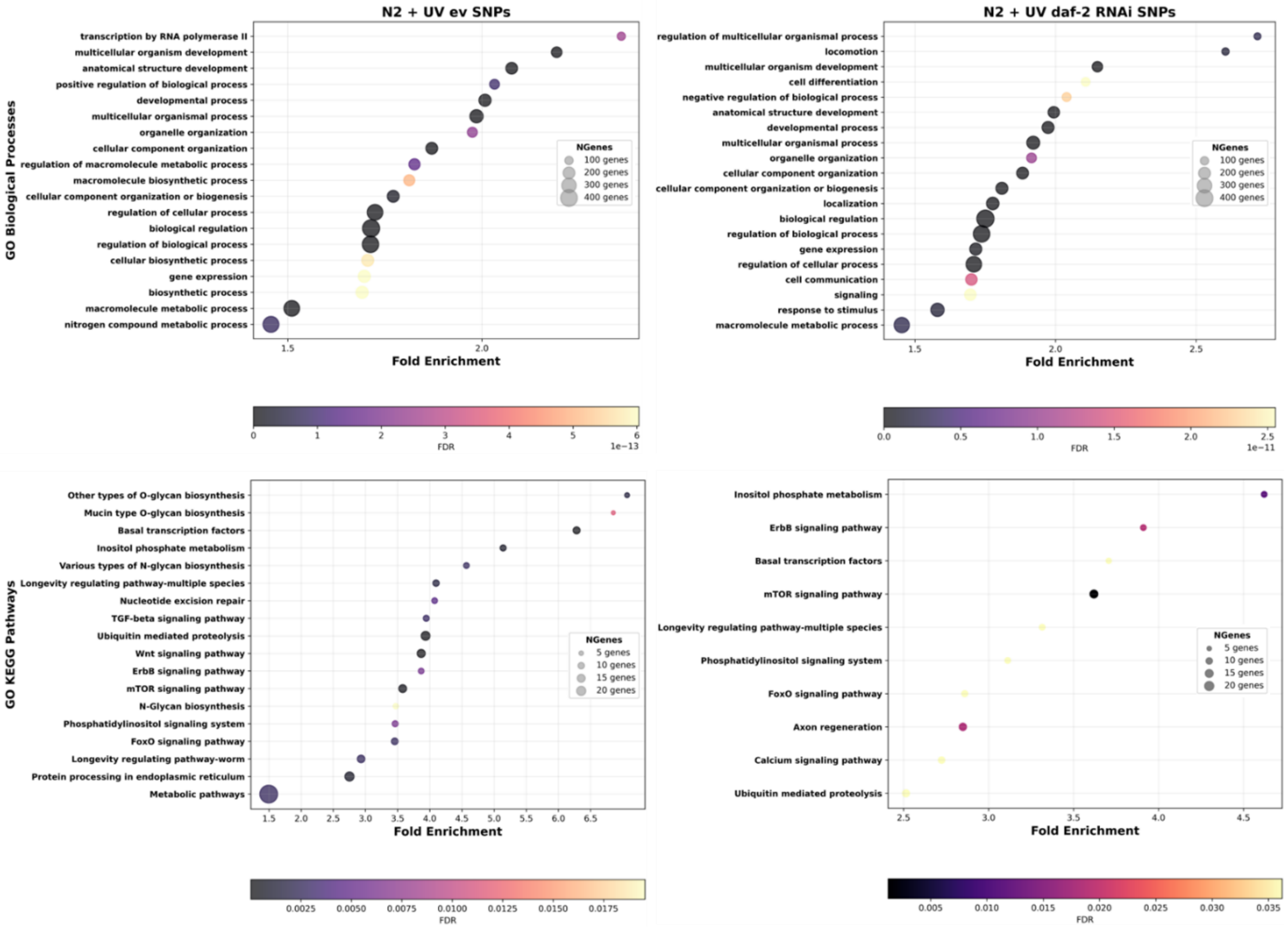


**Supplementary Figure 5.** Functional enrichment analysis of all single nucleotide polymorphism (SNP) mutations in UV-irradiated N2 wild type MA lines under reduced adulthood insulin signalling, via *daf-2* RNAi and in empty vector (ev) controls.

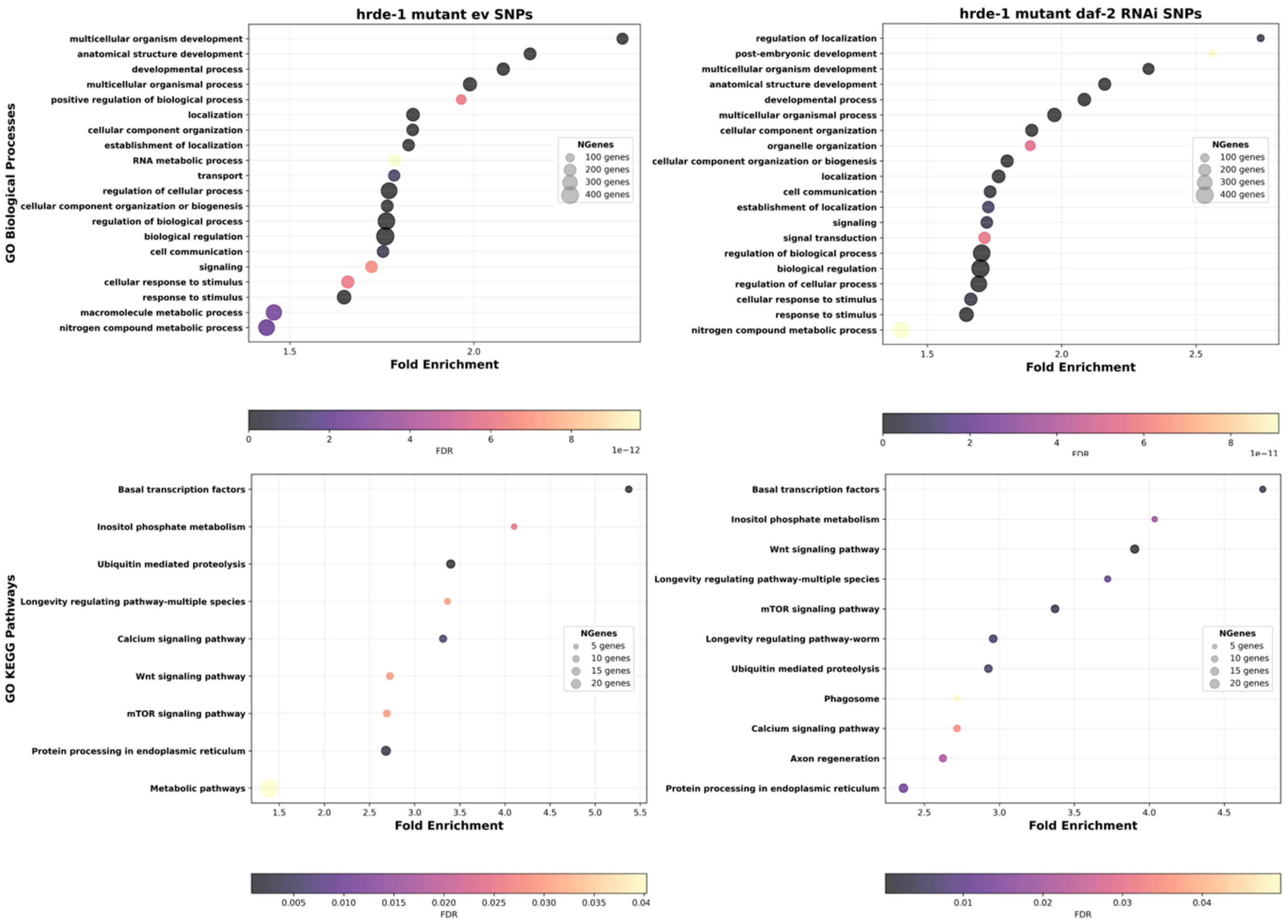


**Supplementary Figure 6.** Functional enrichment analysis of all single nucleotide polymorphism (SNP) mutations in *hrde-1* mutant spontaneous MA lines under reduced adulthood insulin signalling, via *daf-2* RNAi and in empty vector (ev) controls.

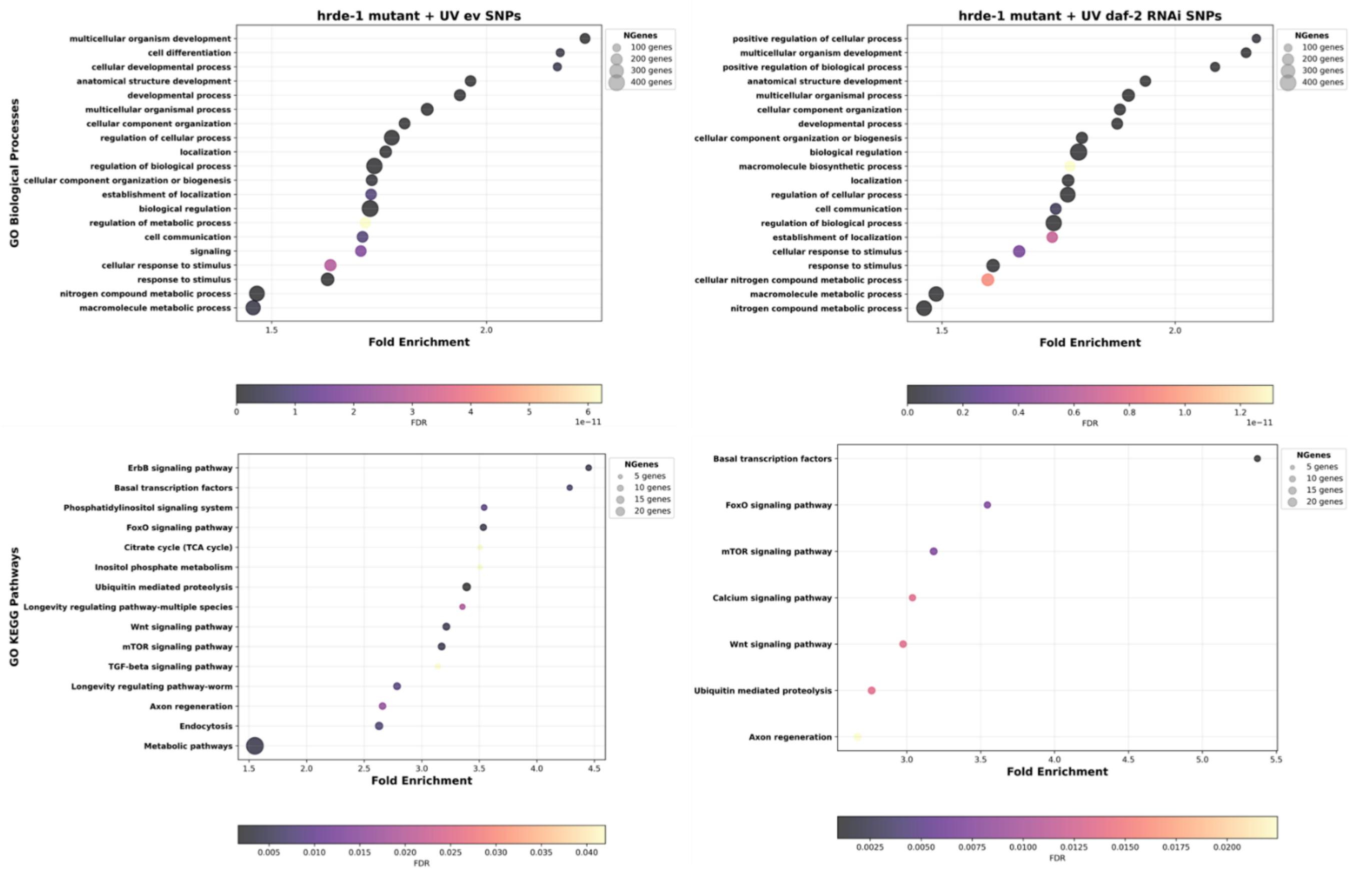


**Supplementary Figure 7.** Functional enrichment analysis of all single nucleotide polymorphism (SNP) mutations in UV-irradiate *hrde-1* mutant MA lines under reduced adulthood insulin signalling, via *daf-2* RNAi and in empty vector (ev) controls.

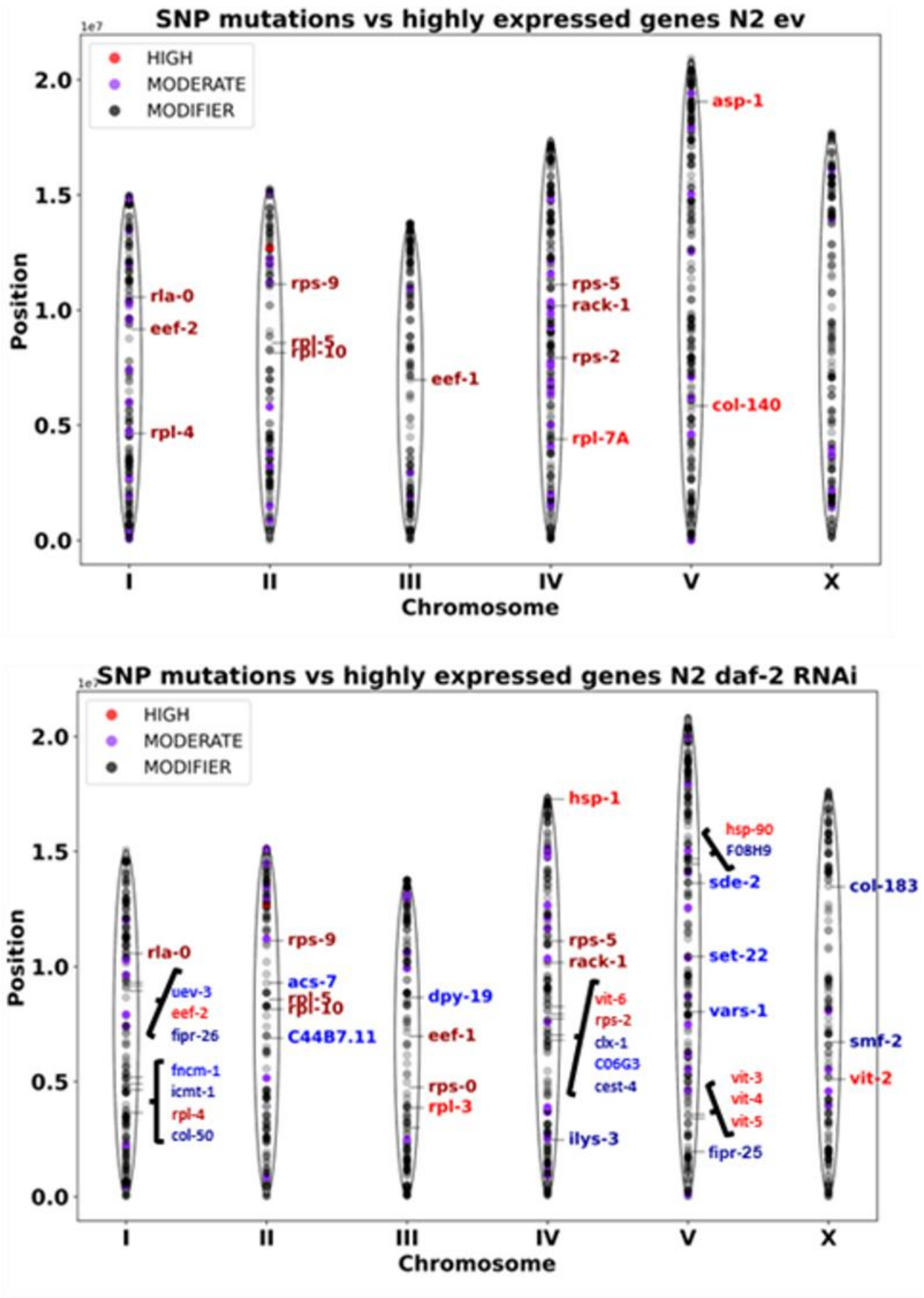


**Supplementary Figure 8. Chromosomal location and impact of N2 spontaneous MA line SNP mutations, and their overlap with highly expressed or upregulated genes from Sultanova et al., 2025.** Genes labelled in the top panel correspond to the top 10 most highly expressed genes in Day 1 (red) or Day 7 (maroon) N2 *C. elegans* worms on the empty vector (ev) control. Several of the same genes overlapped between the two timepoints, resulting in fewer than 20 genes total. Genes in the bottom panel are the top 10 most highly expressed genes in Day 1 (red) or Day 7 (maroon) worms on *daf-2* RNAi, and also the top 10 most highly upregulated genes under *daf-2* RNAi (relative to ev) for Day 1 (mid-blue) and Day 7 (dark blue). Dots indicate all SNP mutations across all MA lines for the respective treatment, colour-coded by the impact of the SNP. Darker shades of grey indicate higher densities of SNPs at a particular genomic location.

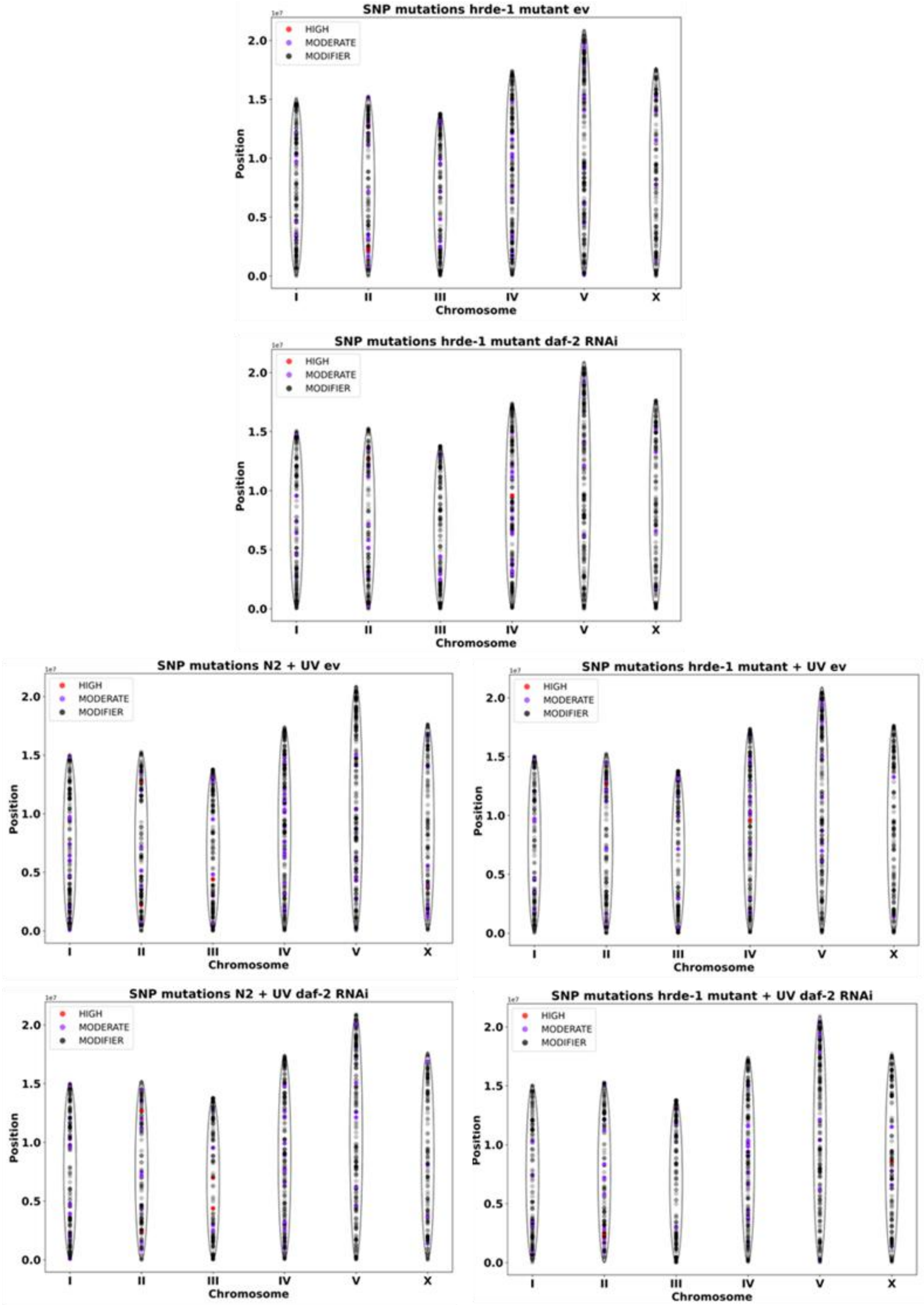


**Supplementary Figure 9. Chromosomal location and impact of N2 and *hrde-1* mutant MA line SNP mutations for empty vector (ev) and *daf-2* RNAi treatments**. Mutations accumulated spontaneously or under UV-induced MA. Dots indicate all SNP mutations across all MA lines for the respective treatment, colour-coded by the impact of the SNP. Darker shades of grey indicate higher densities of SNPs at a particular genomic location. Data for spontaneous MA in N2 background is shown in Suppl. Fig. 8.

#### Supplementary Tables

**Supplementary Table 1. Summary of number of mutations and callable sites for N2 and *hrde-1* mutant across MA lines per treatment.** The full genome size of *C. elegans* is 100.3MB.

| **Treatment** | **Mean (± 1 s.e.) number of mutations** | **Mean number of callable sites** |
|---|---|---|
| **N2 + ev** | 91 (+/-6) | 66986071 |
| **N2 + *daf-2* RNAi** | 80 (+/-4) | 78568333 |
| **N2 + ev + UV** | 144 (+/-13) | 86166818 |
| **N2 + *daf-2* RNAi + UV** | 74 (+/-4) | 87149556 |
| ***hrde-1* mutant + ev** | 60 (+/-4) | 87541840 |
| ***hrde-1* mutant + *daf-2* RNAi** | 59 (+/-4) | 89810292 |
| ***hrde-1* mutant + ev + UV** | 65 (+/-3) | 86303591 |
| ***hrde-1* mutant + *daf-2* RNAi + UV** | 59 (+/-4) | 86650478 |

**Supplementary Table 2.** Statistical output from analysis of SNP mutation rates using a Gaussian GLM.

| | **t** | **p** |
|---|---|---|
| **Full model (SNP mutation rate ~ UV * RNAi * Background):** | | |
| UV x genetic background | -6.595 | **<0.001** |
| RNAi x genetic background | 0.821 | 0.413 |
| UV x RNAi x background | 3.514 | **<0.001** |
| **N2 background:** | | |
| UV | 0.674 | 0.503 |
| RNAi | 1.968 | 0.054 |
| UV x RNAi | 2.362 | **0.022** |
| **Non-irradiated N2 lines:** | | |
| RNAi | 1.987 | 0.057 |
| **UV-irradiated N2 lines:** | | |
| RNAi | 5.153 | **<0.001** |
| ***hrde-1* mutant background:** | | |
| UV | 9.154 | **<0.001** |
| RNAi | 0.155 | 0.877 |
| UV x RNAi | -3.326 | **0.001** |
| **Non-irradiated *hrde-1* mutant lines:** | | |
| RNAi | 0.435 | 0.665 |
| **UV-irradiated *hrde-1* mutant lines:** | | |
| RNAi | -3.189 | **0.003** |

**Supplementary Table 3.** Statistical output from analysis of number of unique SNP mutations per MA line, using a Negative Binomial GLM with log callable sites as the offset, for pairwise comparisons of *daf-2* RNAi versus empty vector treatments. UV-irradiated MA lines did not have matching numbers of MA generations between *daf-2* RNAi and ev treatments, so were excluded from this analysis.

| | z | p |
|---|---|---|
| **Full model (glm.nb(SNPs_unique~RNAi+offset(logcall)):** | | |
| **Non-irradiated N2 lines:** | | |
| RNAi | 2.288 | **0.022** |
| **UV-irradiated N2 lines:** | | |
| RNAi | 5.665 | **<0.001** |
| **Non-irradiated *hrde-1* mutant lines:** | | |
| RNAi | 0.472 | 0.637 |

**Supplementary Table 4.** Full output from Gaussian generalised linear models (GLM) used to analyse transition/tranversion (Ts/Tv) ratios across MA treatments.

| | t | p |
|---|---|---|
| **Full model ( Ts/Tv ~ UV * RNAi * Background):** | | |
| UV x genetic background | -0.593 | 0.554 |
| RNAi x genetic background | -0.743 | 0.459 |
| UV x RNAi x background | 0.988 | 0.325 |
| **N2 background:** | | |
| UV | 0.097 | 0.923 |
| RNAi | -0.154 | 0.878 |
| UV x RNAi | 0.521 | 0.605 |
| **Non-irradiated N2 lines:** | | |
| RNAi | -0.161 | 0.873 |
| **UV-irradiated N2 lines:** | | |
| RNAi | 0.552 | 0.586 |
| ***hrde-1* mutant background:** | | |
| UV | 1.054 | 0.295 |
| RNAi | 1.026 | 0.308 |
| UV x RNAi | -0.940 | 0.350 |
| **Non-irradiated *hrde-1* mutant lines:** | | |
| RNAi | 1.299 | 0.200 |
| **UV-irradiated *hrde-1* mutant lines:** | | |
| RNAi | -0.269 | 0.790 |

**Supplementary Table 5.** Statistical output from Wilcoxon signed rank test of transition/transversion (Ts/Tv) ratios to determine deviance from expected ratio of 0.5. One-sided, paired test was used on data from all MA lines for each treatment separately.

| **Treatment** | **V** | **p** |
|---|---|---|
| **N2 + ev** | 81 | **0.039** |
| **N2 + *daf-2* RNAi** | 101 | **0.009** |
| **N2 + ev + UV** | 66 | **0.001** |
| **N2 + *daf-2* RNAi + UV** | 130 | **0.029** |
| ***hrde-1* mutant + ev** | 288 | **<0.001** |
| ***hrde-1* mutant + *daf-2* RNAi** | 158 | 0.072 |
| ***hrde-1* mutant + ev + UV** | 150 | **0.048** |
| ***hrde-1* mutant + *daf-2* RNAi + UV** | 180 | **0.013** |

**Supplementary Table 6.** Full output from generalised linear model with quasibinomial error structure to account for overdispersion used to analyse fine-scale mutation spectra data (full model:glm(Mutation_type_sum/Total_sum~Treatment,family=quasibinomial,weights=Total_sum)). Overall MA treatment effect (across all eight combinations of RNAi by UV by genetic background) determined for each mutation type using anova likelihood ratio test on the quasibinomial model, then separate pairwise comparisons to determine the effect of *daf-2* RNAi.

| | t | p |
|---|---|---|
| **Mutation type: G->A, C->T** | | |
| MA treatment | | **<0.001** |
| Non-irradiated N2 lines, RNAi: | 0.522 | 0.604 |
| UV-irradiated N2 lines, RNAi: | -0.118 | 0.906 |
| Non-irradiated *hrde-1* mutant lines, RNAi: | -1.781 | 0.075 |
| UV-irradiated *hrde-1* mutant lines, RNAi: | 1.729 | 0.087 |
| **Mutation type: A->G, T->C** | | |
| MA treatment | | 0.594 |
| Non-irradiated N2 lines, RNAi: | 0.245 | 0.807 |
| UV-irradiated N2 lines, RNAi: | -1.961 | 0.054 |
| Non-irradiated *hrde-1* mutant lines, RNAi: | -0.340 | 0.734 |
| UV-irradiated *hrde-1* mutant lines, RNAi: | -0.640 | 0.524 |
| **Mutation type: G->C, C->G** | | |
| MA treatment | | 0.413 |
| Non-irradiated N2 lines, RNAi: | 1.730 | 0.089 |
| UV-irradiated N2 lines, RNAi: | -0.289 | 0.774 |
| Non-irradiated *hrde-1* mutant lines, RNAi: | -0.414 | 0.680 |
| UV-irradiated *hrde-1* mutant lines, RNAi: | -1.894 | 0.062 |
| **Mutation type: G->T, C->A** | | |
| MA treatment | | 0.054 |
| Non-irradiated N2 lines, RNAi: | 1.370 | 0.176 |
| UV-irradiated N2 lines, RNAi: | 1.156 | 0.252 |
| Non-irradiated *hrde-1* mutant lines, RNAi: | 0.565 | 0.573 |
| UV-irradiated *hrde-1* mutant lines, RNAi: | -0.092 | 0.927 |
| **Mutation type: T->G, A->C** | | |
| MA treatment | | **0.033** |
| Non-irradiated N2 lines, RNAi: | -1.613 | 0.112 |
| UV-irradiated N2 lines, RNAi: | 2.120 | **0.038** |
| Non-irradiated *hrde-1* mutant lines, RNAi: | 0.717 | 0.475 |
| UV-irradiated *hrde-1* mutant lines, RNAi: | 1.382 | 0.170 |
| **Mutation type: T->A, A->T** | | |
| MA treatment | | 0.292 |
| Non-irradiated N2 lines, RNAi: | -0.980 | 0.331 |
| UV-irradiated N2 lines, RNAi: | -1.473 | 0.146 |
| Non-irradiated *hrde-1* mutant lines, RNAi: | 0.771 | 0.442 |
| UV-irradiated *hrde-1* mutant lines, RNAi: | -1.829 | 0.071 |

**Supplementary Table 7.** Statistical output from analysis of TE insertion rates using a Gaussian GLM.

| | t | p |
|---|---|---|
| **Full model (TE insertion rate ~ UV * RNAi * Background):** | | |
| UV x genetic background | -5.853 | **<0.001** |
| RNAi x genetic background | -0.526 | 0.599 |
| UV x RNAi x background | 2.899 | **0.004** |
| **N2 background:** | | |
| UV | 0.203 | 0.840 |
| RNAi | -0.272 | 0.786 |
| UV x RNAi | 0.651 | 0.518 |
| **Non-irradiated N2 lines:** | | |
| RNAi | -0.229 | 0.820 |
| **UV-irradiated N2 lines:** | | |
| RNAi | 0.838 | 0.409 |
| ***hrde-1* mutant background:** | | |
| UV | 7.325 | **<0.001** |
| RNAi | 0.628 | 0.531 |
| UV x RNAi | -3.633 | **<0.001** |
| **Non-irradiated *hrde-1* mutant lines:** | | |
| RNAi | 0.981 | 0.331 |
| **UV-irradiated *hrde-1* mutant lines:** | | |
| RNAi | -3.454 | **<0.001** |

**Supplementary Table 8.** Full output from generalised linear model with binomial error structure used to analyse TE classes data. Full model: glm(formula=Number_in_TE_class/Total ~ RNAi * UV * Background, family = binomial,weights = Total).

| | **z** | **p** |
|---|---|---|
| **TE class: DNA** | | |
| UV x genetic background | 3.330 | **<0.001** |
| RNAi x genetic background | 0.901 | 0.367 |
| RNAi x UV | 2.602 | **0.009** |
| UV x RNAi x background | -2.345 | **0.019** |
| Non-irradiated N2 lines, RNAi: | -0.550 | 0.582 |
| UV-irradiated N2 lines, RNAi: | -1.945 | 0.052 |
| Non-irradiated *hrde-1* mutant lines, RNAi: | -2.224 | **0.026** |
| UV-irradiated *hrde-1* mutant lines, RNAi: | 1.371 | 0.170 |
| **TE class: LINE** | | |
| UV x genetic background | -0.475 | 0.635 |
| RNAi x genetic background | -0.595 | 0.552 |
| RNAi x UV | -0.840 | 0.401 |
| UV x RNAi x background | 0.934 | 0.350 |
| Non-irradiated N2 lines, RNAi: | 0.294 | 0.769 |
| UV-irradiated N2 lines, RNAi: | 1.288 | 0.198 |
| Non-irradiated *hrde-1* mutant lines, RNAi: | 1.793 | 0.073 |
| UV-irradiated *hrde-1* mutant lines, RNAi: | 0.682 | 0.495 |
| **TE class: RC** | | |
| UV x genetic background | -3.040 | **0.002** |
| RNAi x genetic background | -0.332 | 0.740 |
| RNAi x UV | -2.162 | **0.031** |
| UV x RNAi x background | 1.376 | 0.169 |
| Non-irradiated N2 lines, RNAi: | 0.741 | 0.459 |
| UV-irradiated N2 lines, RNAi: | 0.848 | 0.397 |
| Non-irradiated *hrde-1* mutant lines, RNAi: | 1.288 | 0.198 |
| UV-irradiated *hrde-1* mutant lines, RNAi: | -1.944 | 0.052 |

**Supplementary Table 9. Number (and %) of SNP effects by impact per treatment.**

| Treatment | High | Moderate | Low | Modifier |
|---|---|---|---|---|
| **N2 + ev** | 3 (0.028%) | 170 (1.585%) | 123 (1.147%) | 10,432 (97.241%) |
| **N2 + *daf-2* RNAi** | 3 (0.031%) | 148 (1.544%) | 89 (0.928%) | 9,348 (97.497%) |
| **N2 + ev + UV** | 6 (0.054%) | 177 (1.580%) | 98 (0.875%) | 10,924 (97.492%) |
| **N2 + *daf-2* RNAi + UV** | 7 (0.075%) | 117 (1.253%) | 86 (0.921%) | 9,130 (97.752%) |
| ***hrde-1* mutant + ev** | 5 (0.049%) | 144 (1.419%) | 97 (0.956%) | 9,905 (97.577%) |
| ***hrde-1* mutant + *daf-2* RNAi** | 4 (0.041%) | 130 (1.348%) | 96 (0.995%) | 9,416 (97.616%) |
| ***hrde-1* mutant + ev + UV** | 4 (0.041%) | 161 (1.652%) | 82 (0.841%) | 9,499 (97.466%) |
| ***hrde-1* mutant + *daf-2* RNAi + UV** | 3 (0.031%) | 149 (1.562%) | 89 (0.933%) | 9,299 (97.474%) |

**Supplementary Table 10. Number of protein-coding SNPs per functional class per treatment.** The ratio of non-synonymous (nonsense and missense) to synonymous (silent) SNPs is also shown.

| Treatment | Nonsense | Missense | Silent | Ratio (non-synonymous / synonymous SNPs) |
|---|---|---|---|---|
| **N2 + ev** | 0 | 170 | 103 | 1.650 |
| **N2 + *daf-2* RNAi** | 0 | 148 | 67 | 2.209 |
| **N2 + ev + UV** | 2 | 177 | 78 | 2.295 |
| **N2 + *daf-2* RNAi + UV** | 3 | 118 | 67 | 1.806 |
| ***hrde-1* mutant + ev** | 0 | 145 | 83 | 1.747 |
| ***hrde-1* mutant + *daf-2* RNAi** | 0 | 131 | 80 | 1.638 |
| ***hrde-1* mutant + ev + UV** | 0 | 162 | 75 | 2.160 |
| ***hrde-1* mutant + *daf-2* RNAi + UV** | 0 | 151 | 83 | 1.819 |

**Supplementary Table 11.** Statistical output from analysis of protein-coding SNP functional classes (missense or silent) using two-tailed z-test with Yates continuity correction (prop.test), involving pairwise comparisons of RNAi treatments and UV treatments.

| Treatment Comparison | | Missense | Silent |
|---|---|---|---|
| **N2: ev vs. *daf-2* RNAi** | $\chi^2$ | 2.004 | 2.004 |
| | df | 1 | 1 |
| | p | 0.157 | 0.157 |
| **N2 + ev: UV vs. no UV** | $\chi^2$ | 2.268 | 2.885 |
| | df | 1 | 1 |
| | p | 0.132 | 0.089 |
| **N2 + UV: ev vs. *daf-2* RNAi** | $\chi^2$ | 1.548 | 1.152 |
| | df | 1 | 1 |
| | p | 0.213 | 0.283 |
| **N2 + *daf-2* RNAi: UV vs. no UV** | $\chi^2$ | 1.388 | 0.715 |
| | df | 1 | 1 |
| | p | 0.238 | 0.400 |
| ***hrde-1* mutant: ev vs. *daf-2* RNAi** | $\chi^2$ | 0.052 | 0.052 |
| | df | 1 | 1 |
| | p | 0.819 | 0.0819 |
| ***hrde-1* mutant + ev: UV vs. no UV** | $\chi^2$ | 0.970 | 0.970 |
| | df | 1 | 1 |
| | p | 0.325 | 0.325 |
| ***hrde-1* mutant + UV: ev vs. *daf-2* RNAi** | $\chi^2$ | 0.611 | 0.611 |
| | df | 1 | 1 |
| | p | 0.435 | 0.435 |
| ***hrde-1* mutant + *daf-2* RNAi: UV vs. no UV** | $\chi^2$ | 0.190 | 0.190 |
| | df | 1 | 1 |
| | p | 0.663 | 0.663 |

**Supplementary Table 12. N2 wild type SNP mutations by fine-scale type across MA treatments.** The most common variant types are highlighted.

| Variants by type | N2 + ev | N2 + *daf-2* RNAi | N2 + ev + UV | N2 + *daf-2* RNAi + UV |
|---|---|---|---|---|
| 3_prime_UTR_variant | 0.54% | 0.43% | 0.46% | 0.55% |
| 5_prime_UTR_premature_start_codon_gain_variant | 0.01% | 0.01% | 0.07% | 0.09% |
| 5_prime_UTR_variant | 0.10% | 0.07% | 0.14% | 0.24% |
| downstream_gene_variant | 39.23% | 39.86% | 38.83% | 39.48% |
| intergenic_region | 6.16% | 6.67% | 6.38% | 6.41% |
| intron_variant | 12.90% | 14.92% | 13.92% | 14.02% |
| missense_variant | 1.58% | 1.54% | 1.58% | 1.25% |
| non_coding_transcript_exon_variant | 0.48% | 0.49% | 0.39% | 0.41% |
| splice_acceptor_variant | 0.03% | 0.03% | 0.03% | 0.03% |
| splice_donor_variant | 0.00% | 0.00% | 0.01% | 0.00% |
| splice_region_variant | 0.13% | 0.13% | 0.07% | 0.06% |
| stop_gained | 0.00% | 0.00% | 0.02% | 0.03% |
| stop_lost | 0.00% | 0.00% | 0.00% | 0.01% |
| synonymous_variant | 1.02% | 0.80% | 0.74% | 0.77% |
| upstream_gene_variant | 37.82% | 35.06% | 37.36% | 36.66% |

**Supplementary Table 13. *hrde-1* mutant SNP mutations by fine-scale type across MA treatments.** The most common variant types are highlighted.

| Variants by type | *hrde-1* mutant + ev | *hrde-1* mutant + *daf-2* RNAi | *hrde-1* mutant + ev + UV | *hrde-1* mutant + *daf-2* RNAi + UV |
|---|---|---|---|---|
| 3_prime_UTR_variant | 0.47% | 0.65% | 0.35% | 0.42% |
| 5_prime_UTR_premature_start_codon_gain_variant | 0.08% | 0.09% | 0.01% | 0.00% |
| 5_prime_UTR_variant | 0.22% | 0.28% | 0.17% | 0.15% |
| downstream_gene_variant | 38.85% | 40.49% | 39.67% | 38.95% |
| intergenic_region | 6.47% | 6.72% | 6.48% | 6.72% |
| intron_variant | 15.38% | 13.47% | 14.97% | 15.10% |
| missense_variant | 1.42% | 1.35% | 1.65% | 1.56% |
| non_coding_transcript_exon_variant | 0.44% | 0.45% | 0.45% | 0.44% |
| splice_acceptor_variant | 0.03% | 0.03% | 0.03% | 0.00% |
| splice_donor_variant | 0.01% | 0.00% | 0.00% | 0.01% |
| splice_region_variant | 0.06% | 0.11% | 0.09% | 0.09% |
| stop_gained | 0.00% | 0.00% | 0.00% | 0.00% |
| stop_lost | 0.01% | 0.01% | 0.01% | 0.02% |
| synonymous_variant | 0.83% | 0.83% | 0.77% | 0.87% |
| upstream_gene_variant | 35.74% | 35.5 | 35.34% | 35.67% |

**Supplementary Table 14.** Statistical output from analysis of frequency of SNP mutations in exons, introns and intergenic regions using two-tailed z-test with Yates continuity correction (prop.test), involving pairwise comparisons of RNAi treatments and UV treatments.

| Treatment Comparison | | Exon | Intergenic | Intron |
|---|---|---|---|---|
| **N2: ev vs. *daf-2* RNAi** | $\chi^2$ | 1.106 | 2.143 | 17.140 |
| | df | 1 | 1 | 1 |
| | p | 0.293 | 0.143 | **<0.001** |
| **N2 + ev: UV vs. no UV** | $\chi^2$ | 2.198 | 0.449 | 4.719 |
| | df | 1 | 1 | 1 |
| | p | 0.138 | 0.503 | **0.030** |
| **N2 + UV: ev vs. *daf-2* RNAi** | $\chi^2$ | 1.230 | 0.002 | 0.060 |
| | df | 1 | 1 | 1 |
| | p | 0.268 | 0.969 | 0.807 |
| **N2 + *daf-2* RNAi: UV vs. no UV** | $\chi^2$ | 2.026 | 0.488 | 2.900 |
| | df | 1 | 1 | 1 |
| | p | 0.155 | 0.485 | 0.089 |
| ***hrde-1* mutant: ev vs. *daf-2* RNAi** | $\chi^2$ | 0.087 | 0.485 | 14.526 |
| | df | 1 | 1 | 1 |
| | p | 0.768 | 0.486 | **<0.001** |
| ***hrde-1* mutant + ev: UV vs. no UV** | $\chi^2$ | 0.620 | <0.001 | 0.649 |
| | df | 1 | 1 | 1 |
| | p | 0.431 | 0.995 | 0.421 |
| ***hrde-1* mutant + UV: ev vs. *daf-2* RNAi** | $\chi^2$ | <0.001 | 0.430 | 0.057 |
| | df | 1 | 1 | 1 |
| | p | 1 | 0.512 | 0.811 |
| ***hrde-1* mutant + *daf-2* RNAi: UV vs. no UV** | $\chi^2$ | 1.312 | 0.104 | 7.125 |
| | df | 1 | 1 | 1 |
| | p | 0.252 | 0.748 | **0.008** |

**Supplementary Table 15. Number (and %) of protein-coding SNPs per chromosome per treatment.** The length of each chromosome is also stated as a percentage of the nuclear genome, to compare the SNP frequency across chromosomes from MA treatments with the relative chromosome length.

| **Treatment** | **I** | **II** | **III** | **IV** | **V** | **X** | **Total** |
|---|---|---|---|---|---|---|---|
| **N2 + ev** | 219 (14.8%) | 194 (13.1%) | 205 (13.9%) | 310 (20.9%) | 326 (22.0%) | 226 (15.3%) | 1480 |
| **N2 + *daf-2* RNAi** | 230 (16.1%) | 201 (14.1%) | 216 (15.1%) | 259 (18.1%) | 315 (22.1%) | 207 (14.5%) | 1428 |
| **N2 + ev + UV** | 238 (15.0%) | 218 (13.8%) | 227 (14.3%) | 339 (21.4%) | 363 (22.9%) | 197 (12.5%) | 1582 |
| **N2 + *daf-2* RNAi + UV** | 218 (16.3%) | 184 (13.7%) | 192 (14.3%) | 270 (20.2%) | 286 (21.4%) | 189 (14.1%) | 1339 |
| ***hrde-1* mutant + ev** | 227 (15.1%) | 213 (14.1%) | 238 (15.8%) | 298 (19.8%) | 320 (21.2%) | 210 (13.9%) | 1506 |
| ***hrde-1* mutant + *daf-2* RNAi** | 194 (13.8%) | 192 (13.6%) | 222 (15.8%) | 285 (20.2%) | 304 (21.6%) | 211 (15.0%) | 1408 |
| ***hrde-1* mutant + ev + UV** | 204 (14.3%) | 199 (13.9%) | 227 (15.9%) | 278 (19.5%) | 303 (21.25) | 216 (15.1%) | 1427 |
| ***hrde-1* mutant + *daf-2* RNAi + UV** | 208 (14.7%) | 197 (13.9%) | 223 (15.8%) | 278 (19.7%) | 303 (21.4%) | 204 (14.4%) | 1413 |
| **Chromosome length as % of nuclear genome** | 15.0% | 15.2% | 13.7% | 17.4% | 20.9% | 17.7% | |

**Supplementary Table 16.** Statistical output from analysis of distribution of SNP mutations across chromosomes relative to chromosomal length using ChiSq goodness of fit test per MA treatment.

| **Treatment** | $\chi^2$ | **df** | **p** |
|---|---|---|---|
| **N2 + ev** | 20.633 | 5 | **<0.001** |
| **N2 + *daf-2* RNAi** | 13.851 | 5 | **0.017** |
| **N2 + ev + UV** | 44.646 | 5 | **<0.001** |
| **N2 + *daf-2* RNAi + UV** | 19.094 | 5 | **0.002** |
| ***hrde-1* mutant + ev** | 22.464 | 5 | **<0.001** |
| ***hrde-1* mutant + *daf-2* RNAi** | 20.399 | 5 | **0.001** |
| ***hrde-1* mutant + ev + UV** | 15.565 | 5 | **0.008** |
| ***hrde-1* mutant + *daf-2* RNAi + UV** | 18.489 | 5 | **0.002** |

**Supplementary Table 17.** Statistical output from analysis of proportion of SNPs in highly expressed genes or highly upregulated genes relative to the proportion of the callable genome that highly expressed or highly upregulated genes occupy respectively. SNP data from non-irradiated N2 MA lines, analysed using two-tailed z-test with Yates continuity correction (prop.test). The top ten most upregulated genes in *daf-2* RNAi treatment relative to ev controls were used, or the top ten most highly expressed genes in the respective treatments. Expression data from Sultanova et al. (2025).

| **Treatment** | | |
|---|---|---|
| **N2 ev SNPs** | $\chi^2$ | 3.759 |
| **(overlap with highly expressed genes)** | df | 1 |
| | p | 0.053 |
| **N2 *daf-2* RNAi SNPs** | $\chi^2$ | 86.276 |
| **(overlap with highly expressed genes)** | df | 1 |
| | p | **<0.001** |
| **N2 *daf-2* RNAi SNPs** | $\chi^2$ | <0.001 |
| **(overlap with highly upregulated genes)** | df | 1 |
| | p | 1 |

#### Supplementary Methods

##### Quality filtering of SNPs

To set the minimum and maximum depth thresholds for SNP variant calling with bcftools mpileup and bcftools filter, depth was calculated as the coverage of each site, summed across all samples. The average depth across all genomic sites and all 165 samples was 39.45X. The sum of depths of all samples (using the average depth across all genomic sites) was

6637.68X. We then set the minimum depth as one sixth of the total summed average coverage per sample (DP=1100), removing sites where more than half of samples had missing data. We set the maximum depth as three times the total summed average coverage per sample (DP=20,000).

**Setting coverage thresholds for mosdepth**

To determine the number of sites accessible for mutation calling with "mosdepth v.0.3.2 " (Pedersen & Quinlan, 2018), we set coverage thresholds using the mean depth per sample (39.45X, from samtools depth). We then calculated minimum and maximum thresholds using the same methods as for the variant calling depth thresholds (above). The minimum coverage was set to one third the mean depth per sample (13X, = 39.45X/3), following the conservative approach of Keane et al. (2013) and the maximum coverage was set to three times the mean depth per sample (118X, = 39.45X x 3) as a conservative approach to avoid false positives due to sequencing errors or in regions where alignment was more difficult (as Fang et al., 2021; Rashid et al., 2022).

**Setting quality thresholds for TE filtering**

Based on recommendations by Keane et al. (2013) we compared several different quality thresholds for filtering TE-induced insertion mutations: FL=6 & GQ>=28; FL=7 & GQ>=20; FL=8 & GQ>=20 and GQ>=1000, FL>=8, where FL was the breakpoint criteria and GQ was the number of supporting reads. We found qualitatively similar patterns of TE numbers between treatments and so used stringent quality filtering on novel TEs of GQ>20; maximum breakpoint criteria, FL=8, to obtain only the highest confidence TEs.

**Testing for an association in the genomic location between highly expressed genes and germline SNPs**

The genomic location of SNPs was displayed on chromosomal ideogram plots. To determine whether the genomic distribution of SNPs across chromosomes was proportional to chromosomal length we used a Chi-squared goodness of fit test. A two-tailed z-test with Yates continuity correction was used to determine whether SNP mutations occurred more or less frequently in highly expressed genes or highly upregulated genes than the proportion of the callable genome that these genes occupied. Gene expression data from Sultanova et al. (2025).

#### Supplementary Results and Discussion

##### Intergenerational effects of *daf-2* RNAi treatment during adulthood

Adulthood-only feeding of *daf-2* RNAi in parents improves fitness of F1 offspring (Dillin et al., 2002; Lind et al., 2019; Duxbury et al., 2022). These F1 fitness benefits can arise through different mechanisms. For example, *daf-2* RNAi results in physiological changes in adults, in the generation that it is applied and can alter maternal provisioning to their eggs and egg size, which contributes to improved F1 generation fitness (Hibshman et al. 2016; Lind et al. 2019). These effects can occur without the direct inheritance of *daf-2* RNAi via the germline to the developing F1. Further, in Duxbury et al. (2022) following 20 generations of UV-induced MA and *daf-2* RNAi feeding in adulthood, the fitness benefits of *daf-2* RNAi persisted even after two generations of common garden rearing (in the absence of *daf-2* RNAi)- suggesting genetic effects of MA under multigenerational *daf-2* RNAi on fitness, rather than direct exposure of offspring to RNAi nor acute effects of UV.

The absence of protective effects of *daf-2* RNAi for germline mutation accumulation that we show here under UV-induced mutagenesis in the *hrde-1* mutant background, are likely due to deficiency in the inheritance of all RNAi-induced silencing, thus limiting germline immortality, rather than the lack of *daf-2* RNAi inheritance per se. It is possible that some *daf-2* RNAi may have been inherited by offspring. However, this would be expected to result in slowed development of F1 progeny and reduced fitness (as Dillin et al., 2002). The effects on F1 development are expected to be weak, since they did not outweigh the improved fitness, reduced extinction and reduced germline mutation rates that we saw in descendants of daf-2 RNAi treated parents.

In summary, any direct inheritance of *daf-2* RNAi is likely to have been weak, did not outweigh the beneficial effects seen in progeny reported in our work and by others, and is not supported by the common garden fitness benefits seen in Duxbury et al 2022.

**Detailed description of gene ontology analysis**

We conducted a gene ontology analysis to explore the most enriched biological processes and pathways for germline SNPs with high, moderate or modifier effects in MA lines uniquely occurring under rIIS relative to the respective controls. Under rIIS, for the N2 spontaneous MA lines, SNPs with modifier effects were enriched for developmental process, metabolism and conserved longevity-regulating pathways, including a significant association with mTOR and TGF-β signalling pathways (Suppl. Figure 4). High or moderate impact SNPs were associated with biological regulation, mRNA surveillance and response to metal ions under rIIS (Suppl. Figure 4). In the respective controls, SNPs with modifier effect were most highly enriched for developmental and reproductive processes, metabolism, proteolysis and mTOR signalling; with high and moderate impact SNPs also associated with development, but also neuronal transport (Suppl. Figure 4). Both treatments had three high impact SNPs in the gene, *tbc-17*, involved in cellular transport and structure, and with a possible role in regulated cell death and tumour immunity. The over-representation of high and moderate impact germline SNP mutations in genes associated with development in N2 spontaneous MA controls, but not in the respective MA lines under rIIS may have contributed to the faster extinction of the former.

Functional annotation of germline SNPs in the N2 UV-induced MA lines under rIIS revealed those with modifier effect were also most highly enriched for developmental processes, metabolism, conserved longevity-regulating pathways including mTOR and FoxO signalling, as well as axon regeneration (Suppl. Figure 5). Under rIIS, high and moderate impact germline SNP mutations were associated with the maintenance of cell junctions, cellular components and biological regulation, including nonsense mutations in membrane gene F31E3.6 and a high impact variant that resulted in the loss of a stop codon in *vab-19,* involved in embryo and vulva development, cytoskeleton organisation and cell junction maintenance (Suppl. Fig. 5). Mutations in human orthologs of *vab-19* are associated with renal cell carcinoma and cerebral palsy (Ding et al., 2003).

In the respective N2 UV-induced MA controls, 'modifier' mutations were most significantly associated with development, biological regulation, metabolism, proteolysis and mTOR signalling; with high and moderate mutations associated with neuronal function (*pals-26*, C33C12.1 and F35A5.1) or cellular transport, structure and potentially regulated cell death and tumour immunity (*tbc-17*) (Suppl. Fig. 5). High impact mutations in *tbc-17* and *pals-26* were also found under rIIS in the N2 UV-induced MA lines, as was enrichment for the FoxO signalling pathway (Suppl. Fig. 5). C33C12.1 and *pals-26* are downstream of insulin signalling pathway genes including *daf-2*, and also *daf-16* and *skn-1* for *pals-26.* C33C12.1 is enriched in motor neurons, *pals-26* in dopaminergic neurons and F35A5.1 in neurofilaments. The high impact variants in F35A5.1 (in controls only) and *pals-26* (in both rIIS and control treatments) were the gain of stop codons that terminated protein synthesis (nonsense mutations). Mutations in human orthologs of F35A5.1 are associated with autoimmune disease of the nervous system and the rare brain disorder, Creutzfeldt-Jakob disease (Culetto & Sattelle, 2000).

Overall, for the N2 MA lines, the most pronounced differences between rIIS and control treatments in the functional enrichment of high and moderate impact germline SNPs, that were most likely to alter protein function and lead to phenotypic effects can be summarised as follows. Under spontaneous MA in controls, there was functional enrichment of high and moderate impact SNPs for developmental processes, whereas under rIIS, only SNPs with modifier effects (likely affecting non-coding regions) were enriched for development and reproduction, so perhaps with reduced detrimental phenotypic effect. Under UV-induced MA, high and moderate impact SNP mutations were generally enriched for a wider variety of critical biological functions, including neuronal function, immunity and embryo and vulval development; with several disease-associated human orthogs. Unique to rIIS were those associated with cellular maintenance, perhaps with beneficial phenotypic effect, whereas those unique to the respective controls had more enrichment for high impact SNPs in genes associated with neuronal function and with human orthologs associated with autoimmune disease and rare brain disorder. The occurrence of a few SNPs in *tbc-17* across all MA treatments was unexpected and the underlying mechanisms are as yet unclear.

We next functionally annotated germline SNPs in the *hrde-1* mutant MA lines to compare patterns of enrichment with the N2 background, between rIIS and control treatments, and between UV-induced and spontaneous MA. In the *hrde-1* mutant spontaneous MA lines, under rIIS, high and moderate impact germline SNPs were enriched for biological regulation and protein processing, and those with modifier impact were significantly enriched for developmental processes, Wnt and mTOR signalling, and longevity-regulating and protein processing pathways (Suppl. Fig. 6). For the respective ev controls, high and moderate SNPs were enriched for cellular maintenance, protein modification, metabolism and dosage compensation; whereas those with modifier effect were enriched for a wider range of functions, including development, regulation, RNA metabolism, protein processing, mTOR signalling and longevity-regulating pathways (Suppl. Fig. 6). In controls relative to the rIIS treatment, the larger range of critical biological functions associated with both high/moderate impact SNPs and those with modifier impact, may have indicated a larger underlying genetic load, that may contribute to detrimental phenotypic effects, if either combined with an additional environmental stress such as UV-irradiation, or after a greater number of spontaneous MA generations. Under rIIS but not in controls, we found a high impact in F27C8.3 that is enriched on excretory canal cells and is affected by genes involved in stress response (*hsp-6* and *aak-2*) and by *clk-1* which is associated with biological timing, growth, development, behaviour, lifespan and ageing.

Finally, we performed functional annotation of germline SNPs in *hrde-1* mutants under UV-induced MA. Under rIIS, high and moderate impact SNPs were significantly associated with development, reproduction, biological regulation, metabolism and cellular maintenance, and those with modifier effect were associated with biosynthesis and transport, in addition to development and regulation, with highest enrichment for FoxO and mTOR signalling pathways (Suppl. Fig. 7). Under rIIS, the three high impact variants resulted in the loss of a stop codon, and occurred in motor neuron gene, CC33C12.1, and *vab-19*, associated with embryo and vulval development and cellular maintenance, and in *sdha-1*, which is associated with mitochondrial electron transport. Mutations in the human ortholog of *sdha-1* are associated with the rare neurodegenerative Leigh syndrome.

The strong enrichment for SNPs in genes with potentially detrimental effects is likely to have contributed to the rapid extinction seen under UV-induced MA and rIIS in *hrde-1* mutant lines. In respective *hrde-1* mutant UV-induced MA control lines, germline SNPs with modifier effect were enriched for development, regulation and metabolism, including proteolysis, FoxO, Wnt, mTOR and metabolic pathways (Suppl. Fig 7), but there was no significant functional enrichment of high and moderate impact SNPs unique to these control lines, perhaps indicative of reduced phenotypic impact.

**Further exploratory analyses of the genomic location of germline SNPs**

SNPs occurred less frequently on the X chromosome (12-15% of SNPs) relative to the proportion of the genome it contributes (17.7%), and also less frequently on chromosome II (13-14%) relative to 15.2%, but more frequently on all other autosomes than their relative contribution to the genome (Suppl. Tables 15 & 16). Across MA treatments, high and moderate impact SNPs often occurred outside of the genomic locations where there were high densities of modifier SNPs (Suppl. Fig.s 8 & 9).

For N2 non-irradiated MA treatments, we overlapped the genomic location of SNP mutations with a separate *C. elegans* whole-transcriptome RNA-seq dataset (Sultanova et al., 2025) that provided an extensive catalogue of gene expression in Day 1 and Day 7 N2 worms on ev control or adulthood *daf-2* RNAi treatments in a single generation, in the absence of MA or UV-irradiation. We aimed to test the hypothesis that there may be fewer mutations in regions of the genome where there are more highly expressed genes that are more likely to be associated with essential functions and therefore perhaps subject to more transcription-coupled repair (Monroe et al., 2022; 2023; but see Wang et al., 2023). As such, we first overlapped the top 10 most highly expressed genes from Day 1 and from Day 7 worms in respective ev or *daf-2* RNAi treatments (from Sultanova et al., 2025) with the SNP mutations for that corresponding treatment. In many cases, several of the same 10 highly expressed genes overlapped between Day 1 and Day 7 timepoints, resulting in fewer than 20 genes total from the two timepoints. We then investigated whether the top 10 genes that are most highly upregulated under *daf-2* RNAi relative to ev in Day 1 or Day 7 N2 worms (from Sultanova et al., 2025), and thus may also be subject to elevated levels of transcription-coupled repair, were located in regions of the genome with fewer SNP mutations for our N2 non-irradiated MA lines on *daf-2* RNAi.

SNP mutations for non-irradiated N2 ev control MA lines did not occur significantly more or less frequently in the highly expressed genes than expected- relative to proportion of the callable genome that these genes occupy (Suppl. Table 17). In fact, three out of the 13 most highly expressed genes (*rpl-4, rack-1* and proteolytic *asp-1*; from Day 1 and Day 7 expression data combined; Sultanova et al., 2025), overlapped with genomic regions with moderate impact SNP mutations (Suppl. Fig. 8), despite predictions of higher transcription-coupled repair. For N2 worms on adulthood *daf-2* RNAi, SNP mutations occurred significantly more frequently in the most highly expressed genes, than would be expected for the relative proportion of the callable genome that these genes occupy (Suppl. Table 17). Additionally, two out of the 19 most highly expressed genes (ribosomal *rps-9* and *rack-1*; from Days 1 and 7 combined) overlapped with moderate impact SNPs (Suppl. Fig. 8). This is contrary to the prediction of fewer mutations in more highly expressed genes (Monroe et al., 2022; 2023).

Finally, SNP mutations for non-irradiated N2 MA lines on *daf-2* RNAi did not occur significantly more or less frequently in the highly expressed genes than expected- relative to proportion of the callable genome that these genes occupy (Suppl. Table 17). In fact, two out of the 20 most highly upregulated genes overlapped with regions of high impact SNP mutations (*clx-1*, associated with collagen, and *ilys-3*, associated with innate immune response; Suppl. Fig. 8).

Overall, there was not conclusive support for the hypothesis that SNP mutations occurred outside of genomic regions where the most highly expressed or upregulated genes were located. It is possible instead that certain more highly transcribed regions were more vulnerable to mutations, as seen in several organisms (Kim & Jinks-Robertson, 2012), thus masking an overall genome-wide correlation between gene expression and SNP mutations. Further, it is possible that other factors such as recombination rate, which is correlated with mutation rate across several species (Cutter & Choi, 2010; Arbeithuber et al., 2015; Smith et al., 2018) may have had more impact on the genomic location of mutations. Indeed, in *C.*

*elegans*, the chromosomal arms have higher recombination and an increased incidence of repetitive elements, but lower gene densities than the centre and tips of their chromosomes (Rockman & Kruglyak, 2009). Across treatments we also found an overall lower incidence of SNP germline mutations in the chromosome centres, but no apparent decrease in SNPs at the chromosome tips (Suppl. Fig. 8 & 9).

#### Supplementary References


Arbeithuber B, Betancourt AJ, Ebner T, Tiemann-Boege I. Crossovers are associated with mutation and biased gene conversion at recombination hotspots. Proc Natl Acad Sci U S A. 2015 Feb 17;112(7):2109-14. doi: 10.1073/pnas.1416622112.

Culetto E., David B. Sattelle, A role for *Caenorhabditis elegans* in understanding the function and interactions of human disease genes, *Human Molecular Genetics*, Volume 9, Issue 6, 2000, Pages 869–877.

Cutter AD, Choi JY. Natural selection shapes nucleotide polymorphism across the genome of the nematode Caenorhabditis briggsae. Genome Res. 2010 Aug;20(8):1103-11. doi: 10.1101/gr.104331.109.

Ding M, Goncharov A, Jin Y, Chisholm AD. C. elegans ankyrin repeat protein VAB-19 is a component of epidermal attachment structures and is essential for epidermal morphogenesis. Development, 2003;130(23):5791-5801.

Fang, L.T., Zhu, B., Zhao, Y. *et al.* Establishing community reference samples, data and call sets for benchmarking cancer mutation detection using whole-genome sequencing. *Nat Biotechnol* **39**, 1151–1160 (2021).

Hibshman, J.D., Hung, A. & Baugh, L.R. 2016. Maternal diet and insulin like signaling control intergenerational plasticity of progeny size and starvation resistance. PLoS Genet., 14, e1007639.

Kim N, Jinks-Robertson S. Transcription as a source of genome instability. Nat Rev Genet. 2012 Feb 14;13(3):204-14. doi: 10.1038/nrg3152

Monroe, J.G., Srikant, T., Carbonell-Bejerano, P. *et al.* Mutation bias reflects natural selection in *Arabidopsis thaliana*. *Nature* 602, 101–105 (2022).

Monroe, J.G., Murray, K.D., Xian, W. *et al.* Reply to: Re-evaluating evidence for adaptive mutation rate variation. *Nature* 619, E57–E60 (2023).

Rashid, I., Campos, M., Collier, T. *et al.* Spontaneous mutation rate estimates for the principal malaria vectors *Anopheles coluzzii* and *Anopheles stephensi*. *Sci Rep* **12**, 226 (2022).

Rockman MV, Kruglyak L. Recombinational landscape and population genomics of Caenorhabditis elegans. PLoS Genet. 2009 Mar;5(3):e1000419. doi: 10.1371/journal.pgen.1000419.

Smith TCA, Arndt PF, Eyre-Walker A. Large scale variation in the rate of germ-line de novo mutation, base composition, divergence and diversity in humans. PLoS Genet. 2018 Mar 28;14(3):e1007254. doi: 10.1371/journal.pgen.1007254.

Wang, L., Ho, A.T., Hurst, L.D. *et al.* Re-evaluating evidence for adaptive mutation rate variation. *Nature* 619, E52–E56 (2023).
